# A novel downstream factor in willows replaces the ancestral sex determining gene

**DOI:** 10.1101/2024.10.14.618180

**Authors:** Yuàn Wang, Zhi-Qing Xue, Ren-Gang Zhang, Elvira Hörandl, Xiao-Ru Wang, Judith E. Mank, Li He

**Affiliations:** Eastern China Conservation Centre for Wild Endangered Plant Resources, Shanghai Chenshan Botanical Garden, Shanghai, 201602, China; Yunnan Key Laboratory for Integrative Conservation of Plant Species with Extremely Small Populations, Kunming Institute of Botany, Chinese Academy of Sciences, Kunming 650201, Yunnan, China; Department of Systematics, Biodiversity and Evolution of Plants (with Herbarium), University of Goettingen, Göttingen, Germany; Department of Ecology and Environmental Science, Umeå Plant Science Centre, Umeå University, Umeå, Sweden; Department of Zoology and Biodiversity Research Centre, University of British Columbia, Vancouver, BC, Canada

**Author notes:** Corresponding Author: Li He.

**Keywords:** dioecy, polyploidy, sex-determining gene turnover, sex chromosomes

## Abstract

Sex chromosome turnover has occurred in many groups, and is mediated by the translocation or duplication of apical sex-determining genes, or the replacement of original sex determination genes by new ones. In dioecious plants, the former frequently occurs, while the latter is rarely reported. Here, we assembled four haplotype-resolved chromosome-level genomes from four lineages of the *Salix polyclona* complex, including three diploids (E, E-TS, and W1) and one autotetraploid (W2). Our analyses reveal that diploids have a ZW/ZZ system, while autotetraploid has a ZZZW/ZZZZ system on chromosome 15. The apical sex-determining genes of Salicaceae, *ARR17*-like duplicates on chromosomes 15 and 19, which can indirectly turn-on/off the expression of *PISTILLATA* (*PI*) gene and determine sex, appears to have lost the function of sex determination in the *S. polyclona* complex. We found a novel sex determining factor on the 15W of the complex, namely partial *PI*-like duplicates, which has taken over the function from the *ARR17*-like duplicates. In conclusion, a new dominant sex-determining gene was recruited in the *S. polyclona* complex, replacing the ancestral apical sex-determining *ARR17*-like duplicates. The newly identified partial *PI*-like duplicates exhibit a direct influence on the downstream intact *PI*-like duplications, providing valuable insights into the evolutionary trajectory of the top-down sex determination pathway.

## Introduction

Many recent studies have revealed extensive changes in the location of sex chromosomes ^1–5^, much of which is thought to be due to changes in the location of the apical sex determination gene or rewiring of the sex determination pathway. Sex chromosome turnover can occur when apical sex-determining genes move from one chromosome to another ^6–9^, as has occurred in the Salicaceae when translocations and duplications of the *ARABIDOPSIS RESPONSE REGULATOR 17* (*ARR17*)*-*like sequence, the proposed ancestral sex determination gene of *Populus* and *Salix* ^10–17^ induced changes in the chromosomes associated with sex ^11,13^. In *Populus*, *ARR17* fragments in the Y-specific region produce small RNAs that silence intact *ARR17*-like genes, thereby determining males ^10^. In the sister genus *Salix*, partial *ARR17*-like duplicates have been also found in sex-linked regions (SLRs) of both the *Salix*-clade, which exhibits an XY system on chromosome 7, and the *Vetrix*-clade, which includes both XY and ZW systems on chromosome 15 ^11,13,15,18^. The intact *ARR17*-like duplicates indirectly suppress the B-class gene *PISTILLATA* (*PI*), which is essential for stamen development ^19^. Similarly, *PI*-like genes also participate the downstream regulation of male sex determination in persimmon ^20^.

The genetic pathway of sex determination has rarely been discussed in dioecious plants ^21^, and even less is known about how sex determination network rewiring might affect plant sex chromosome turnover. Wilkins proposed that novel sex-determination genes evolve from the top of the cascade, with genes taking control of the sex determination pathway at the initial stages ^22^. The *SRY* gene has been identified as the primary sex-determining gene in mammals ^23^, playing a crucial role in the upstream of sex determination pathway ^24^. Meanwhile, downstream factors of *SRY* gene such as *DMRT1* genes or homologs, *amh*, and *Gsdf^Y^* are often recruited as key sex-determining genes in non-mammalian vertebrates ^25–33^. In *Drosophila*, upstream elements *Sxl* of the sex-determination pathway were recruited to regulate sex ^34^. We are interested in exploring whether dioecious plants exhibit a pattern of sex determination pathway redeployment similar to that in animals, and in understanding the evolutionary process of this sex-determination pathway within dioecious plants.

The genus *Salix* is an emerging model of plant sex chromosome evolution due to the variability described above. *Salix polyclona* is a diploid-autotetraploid complex of *Salix* genus *Vetrix*-clade. Its diploid and autotetraploid lineages have female heterogametic sex chromosomes ^35^. Species could have variable non-recombining SLRs in different populations ^36^. To explore the changes of sex chromosomes, SLRs, and sex determining genes in closely related lineages, we sequenced and assembled four lineages representing the core phylogenetic groups of the *S. polyclona* complex ^35^, including one autotetraploid, West 2 (W2), and three diploid individuals, West 1 (W1), East (E), and Taishan population of East (E-TS). Using our haplotype-resolved chromosome-level assembled genomes of the four lineages and re-sequencing datasets, we identified the sex chromosomes and SLRs in each. We determined the old apical and novel downstream sex-determining genes and their evolutionary trajectories in the *S. polyclona* complex, which we then validated with our expression data. Our results show that members within the *S. polyclona* complex have recruited new sex-determining factors, without changing sex chromosomes.

## Results

### Chromosome-level genome assemblies for *Salix polyclona*

PacBio HiFi reads and Illumina short reads were used to generate genome assemblies for diploid *S. polyclona* samples from the E, E-TS, and W1 lineages. We produced ∼46 Gb (>100×) PacBio HiFi reads and ∼42 Gb (>80×) Illumina short reads for *S. polyclona*-E and *S. polyclona*-W1, and ∼27 Gb (∼52×) PacBio HiFi reads and ∼35 Gb (∼70×) Illumina short reads for *S. polyclona*-E-TS (Supplementary Table 1). In addition to ∼43 Gb (∼56×) PacBio HiFi reads and ∼110 Gb (∼124×) Illumina short reads, ∼54 Gb (∼72×) Oxford Nanopore Technology (ONT) long reads were used to assemble tetraploid *S. polyclona*-W2 (Supplementary Table 1). Primary assemblies were improved to chromosome-level with 87–136× Hi-C reads. After primary assembly, comparison, correction, polishing, and scaffolding, we obtained haplotype-resolved chromosome-level genomes for the four lineages of *S. polyclona,* including a tetraploid and three diploids (Supplementary Figs. 1-4).

The genome size of the three diploids was 785–831 Mb, with 38 pseudochromosomes including two haplotypes (*a* and *b*) (Fig. 1(a)). The tetraploid *S. polyclona*-W2 assembly was 1,507 Mb, with a final chromosome-scale assembly of 77 pseudochromosomes, including four haplotypes (a, b, c and d), and a potential B chromosome, chromosome 20 (Fig. 1(a), Supplementary Fig. 4). We obtained four high-quality genome assemblies of *S. polyclona* (E, E-TS, W1, and W2), and the contig N50 length was 19–23 Mb (Table 1). In addition, both of the HiFi reads and Illumina short reads map to more than 99% of assembled genomic regions. The completeness of genomes was demonstrated by the Benchmarking Universal Single-Copy Orthologs (BUSCO) assessment, suggesting the completeness of four genomes is more than 98% (Supplementary Table 2).

**Figure 1.**
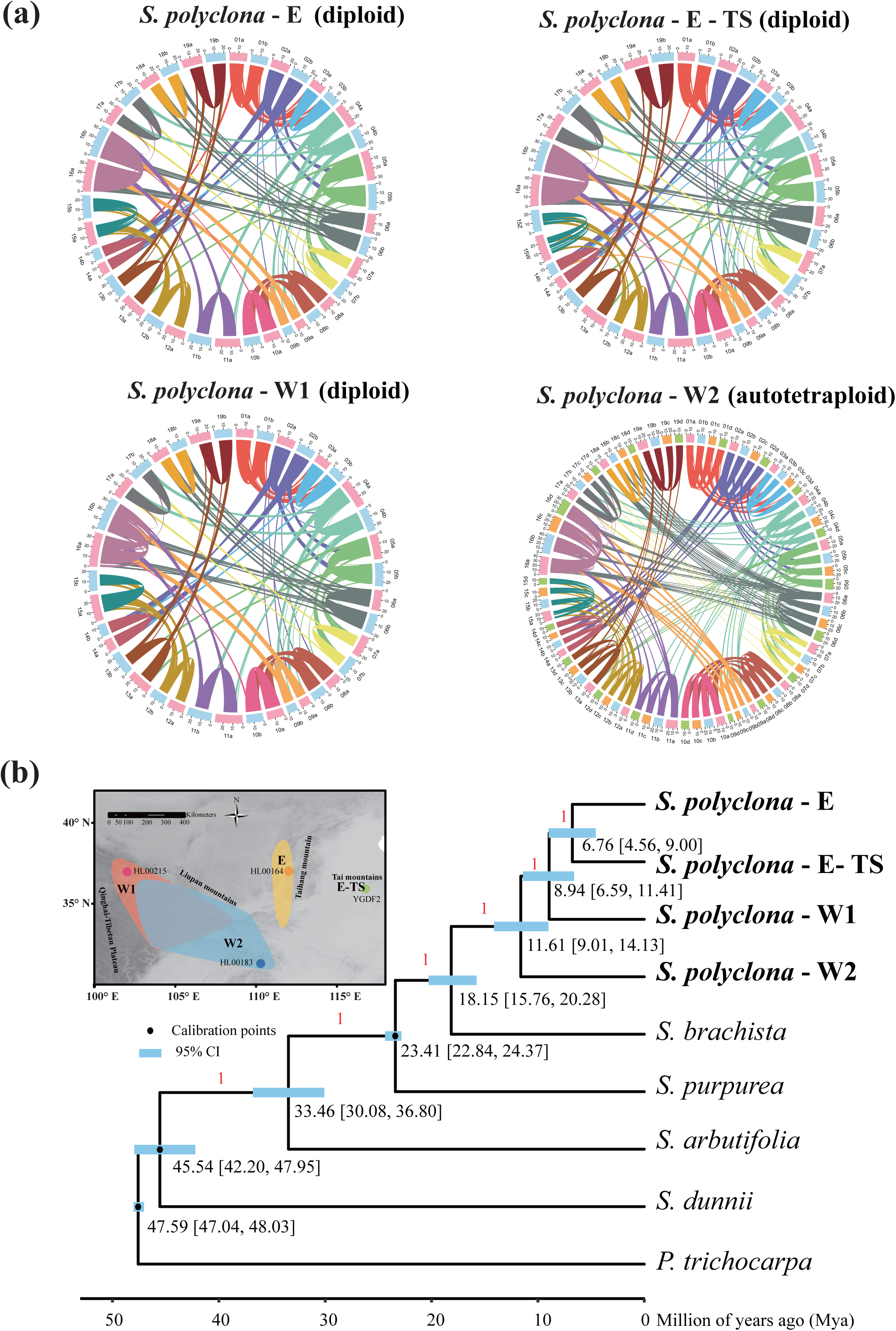
Genome synteny and phylogenetic relationship of the *Salix polyclona* complex. (a) Genome synteny of the *S. polyclona* complex (E, E-TS, W1 and W2). (b) Distribution and divergence times of the *S. polyclona* complex. The map presents the distribution area of the sequenced species, adapted from He et al. ^35^. Inferred phylogenetic tree and divergence times based on genome sequences of eight willows and the outgroup *P. trichocarpa*. Blue node bars are 95% confidence intervals (CI), and the black nodes are three fossil calibration points ^77^. Red numbers marked above branches represent support values. Black numbers marked around nodes represent divergence time.

**Table 1.**
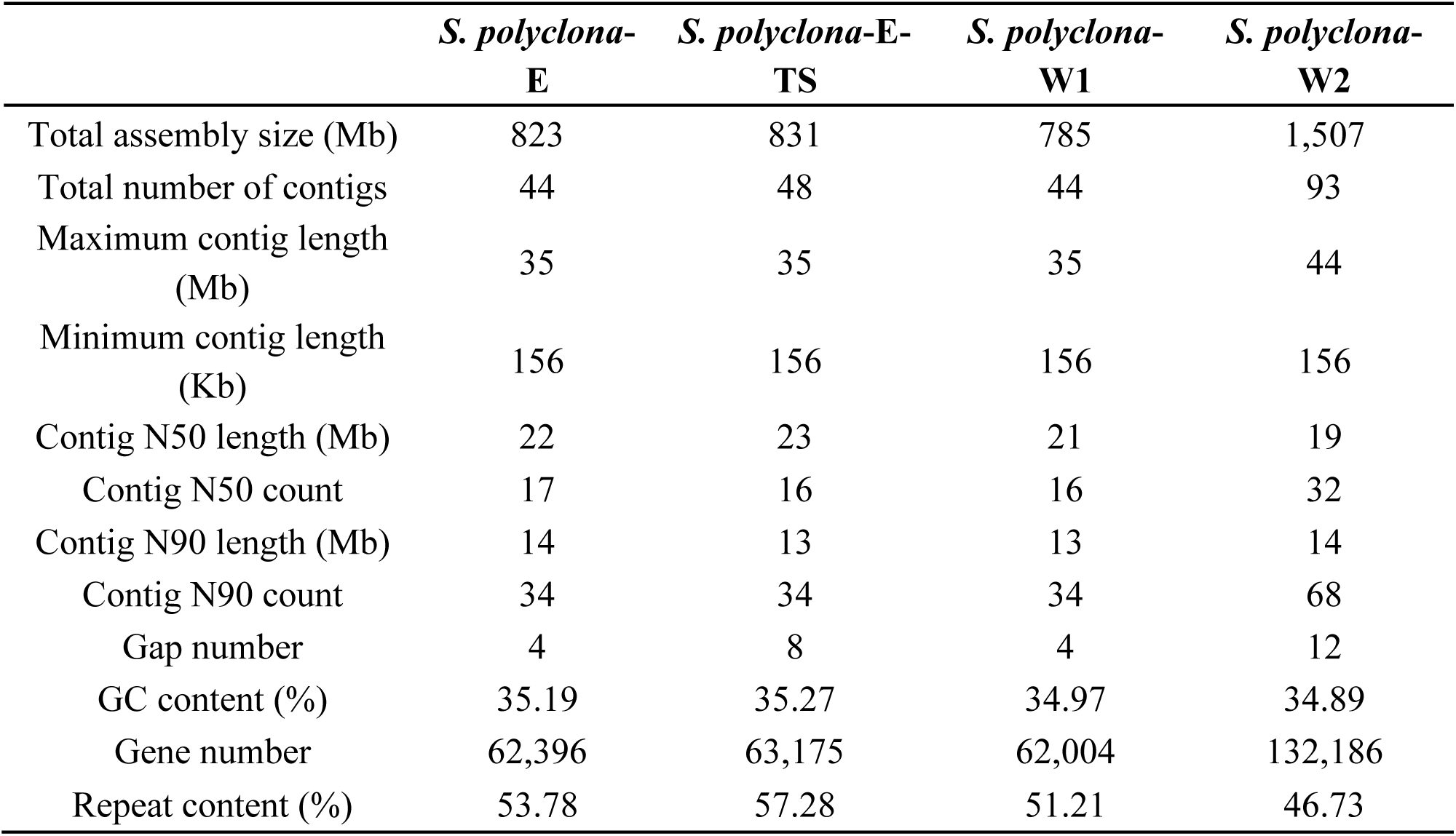
Statistics of the *Salix polyclona* genome assemblies.

|  | *S. polyclona*-<br>E | *S. polyclona*-E-TS | *S. polyclona*-<br>W1 | *S. polyclona*-<br>W2 |
| --- | --- | --- | --- | --- |
| Total assembly size (Mb) | 823 | 831 | 785 | 1,507 |
| Total number of contigs | 44 | 48 | 44 | 93 |
| Maximum contig length<br>(Mb) | 35 | 35 | 35 | 44 |
| Minimum contig length<br>(Kb) | 156 | 156 | 156 | 156 |
| Contig N50 length (Mb) | 22 | 23 | 21 | 19 |
| Contig N50 count | 17 | 16 | 16 | 32 |
| Contig N90 length (Mb) | 14 | 13 | 13 | 14 |
| Contig N90 count | 34 | 34 | 34 | 68 |
| Gap number | 4 | 8 | 4 | 12 |
| GC content (%) | 35.19 | 35.27 | 34.97 | 34.89 |
| Gene number | 62,396 | 63,175 | 62,004 | 132,186 |
| Repeat content (%) | 53.78 | 57.28 | 51.21 | 46.73 |

We predicted 62,396, 63,175, 62,004 and 132,186 genes for whole genomes of *S. polyclona*-E, *S. polyclona*-E-TS, *S. polyclona*-W1 and *S. polyclona*-W2, including 58,669, 58,623, 58,703, 122,173 protein-coding genes, respectively (Supplementary Table 3). For protein-coding genes, the average gene region length ranges from 3614.8 bp to 3753.2 bp, and the average coding sequence length ranges from 1328.1 bp to 1348.9 bp. Among the predicted protein-coding genes, ∼97.5% of them could be annotated by at least one of the annotation methods, which includes eggNOG-mapper, DIAMOND, and InterProScan (Extended Data Table 1). There are 442.83 Mb (53.78%), 476 Mb (57.28%), 401.95 Mb (51.21%), and 704.12 Mb (46.73%) repetitive sequences in genomes of *S. polyclona*-E, *S. polyclona*-E-TS, *S. polyclona*-W1 and *S. polyclona*-W2, respectively (Extended Data Table 2).

### Ploidy and phylogenetic analysis

We used flow cytometry analysis to predict ploidy, and our results suggest that the E, E-TS and W1 lineages of *S. polyclona* are diploids, while W2 is tetraploid (Supplementary Fig. 5). Our *k-mer* results confirmed the latter is indeed autotetraploid, which is consistent with previous result ^35^ (Supplementary Fig. 6).

In order to determine phylogenetic position of *S. polyclona* complex, we obtained 7,230 single-copy genes to reconstruct the species tree of the *S. polyclona* complex and related species, using *Populus trichocarpa* as an outgroup (Supplementary Table 4). We estimate that the split between the *S. polyclona* complex and *S. brachista* occurred 18.15 million years ago (Mya), and the autotetraploid *S. polyclona*-W2 and diploids *S. polyclona* (E, E-TS and W1) diverged 11.61 Mya (Fig. 1(b)), consistent with previous appraisals ^35^.

### Identification of SLRs within the *S. polyclona* complex

Chromosome quotient (CQ), *F*_ST_, synteny and inversion analyses were used to determine the SLRs of *S. polyclona* complex. We obtained 61-83 million clean Illumina reads per individual (mean 66) of the newly sequenced 41 diploids, and 253-263 million reads per individual (mean 256) of the newly sequenced nine tetraploids. The average read depths of the diploids ranged from 19.3× to 28.7×, and the tetraploids ranged from 80.8× to 87.2× using the relevant lineages as the reference genome. Our M:F (male:female) CQ ^37^ analysis confirmed that all four lineages of *S. polyclona* have a female heterogametic sex chromosomes on chromosome 15 (Fig. 2(a)), and we detected the SLRs’ range based on CQ results using changepoint analysis, which covered by the final SLRs’ boundaries (Fig. 2(a)(b), Supplementary Fig. 7(a)(b), Supplementary Fig. 8(a)(b), Supplementary Fig. 9(a)(b), Supplementary Fig. 10(a)(b), Supplementary Table 5).

**Figure 2.**
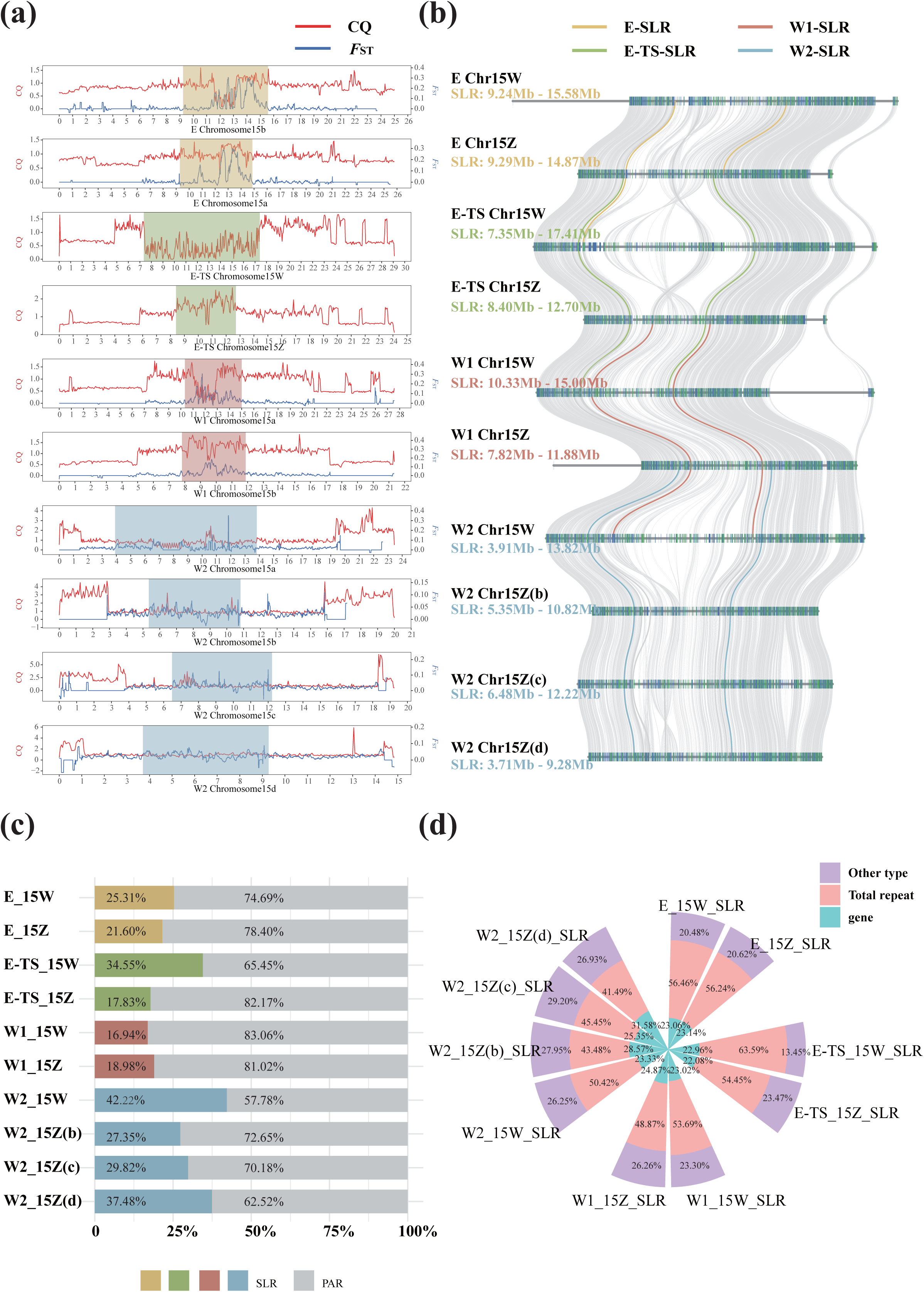
Sex-linked regions in the *S. polyclona* complex (E, E-TS, W1, and W2). (a) The CQ (male vs. female alignment) and *F*_ST_ results on chromosome 15s. The colored regions on lines represent SLRs as determined by CQ, *F*_ST_ and synteny analysis, see Methods. (b) The synteny results between sex chromosomes. Different colored lines mark the sex-linked regions (SLRs). (c) The length percent of SLRs and PARs. (d) The genomic composition in SLRs.

We obtained 4,820,482 and 4,843,879 SNPs from 35 individuals (17 females and 19 males, 17 of them newly sequenced) for haplotypes *a* (including 15Z/15a) and *b* (including 15W/15b) of E, 5,598,460 and 5,609,947 SNPs from 36 individuals (18 females and 18 males, 14 of them newly sequenced for this study) for haplotypes *a* (including 15W/15a) and *b* (including 15Z/15b) of W1, 8,550,694 and 8,312,008 SNPs from 18 individuals (nine females and nine males, nine newly sequenced) for haplotypes *a* & *b* (including 15W(a) and 15Z(b)) and haplotypes *c* & *d* (including 15Z(c) and 15Z(d)) of W2, and used these to calculate the *F*_ST_ values between the male and female genomes (Fig. 2(a), Supplementary Fig. 7(c)(d), Supplementary Fig. 9(c)(d), Supplementary Fig. 10(c)∼(f)). The SLRs identified by *F*_ST_ is broadly consistent with CQ identified regions (Supplementary Table 5).

Synteny analysis among sex chromosomes was also used to identify the SLRs. Inversions were covered by SLRs (Fig. 2(b), Supplementary Fig. 7(e), Supplementary Fig. 8(c), Supplementary Fig. 9(e), Supplementary Fig. 10(g)). So, the final SLR determination combined the results of CQ, *F*_ST_, location of inversions, and syntenic relationships between the identified regions.

The 15W-SLR of *S. polyclona* ranges from 9.24Mb – 15.58Mb, 7.35Mb – 17.41Mb, 10.33Mb – 15.00Mb, 3.91Mb – 13.82Mb on E, E-TS, W1 and W2, and their corresponding 15Z-SLR ranges from 9.29Mb – 14.87Mb, 8.40Mb – 12.70Mb, 7.82Mb – 11.88Mb, 5.35Mb – 10.82Mb (Z(b)), 6.48Mb – 12.22Mb (Z(c)), 3.71Mb – 9.28Mb (Z(d)), respectively (Supplementary Table 5, Fig. 2(b)). For autotetraploid *S. polyclona*-W2, although the 15b and 15d don’t show obvious CQ features of the Z chromosome, we found that the six ancient sex-linked genes on chromosomes 15b, 15c and 15d clustered with Z gametologs, while the homologous genes on chromosome 15a clustered with W gametologs, and partial *ARR17*-like duplicates on chromosomes 15b, 15c, 15d (refer to section ‘The evolution of SLRs in four lineages of *S. polyclona*’ for more details). These results suggest that the *S. polyclona*-W2 has a ZZZW/ZZZZ sex determination system, consistent with He et al.’s hypothesis ^35^.

The proportion of the W-linked region of E-TS (34.55%) and W2 (42.22%) is significantly higher than in E (25.31%) and W1 (16.94%) (Fig. 2(c)), which may be related to the sex chromosome turnover/recombination of E and W1, as detailed in the next section. In addition, there are more genes and duplicates in W-linked region of E-TS (528 genes, 147 duplicates) and W2 (586 genes, 116 duplicates) than in E (401 genes, 69 duplicates) and W1 (314 genes, 79 duplicates) (Supplementary Tables 6-8, Extended Data Tables 3-4). The percentage of repeated sequences within the 15W-SLR of the *S. polyclona* complex is higher than the average in the whole genome (Fig. 2(d), Extended Data Table 2), consistent with non-recombining regions in other willows ^12,15^.

### The evolution of SLRs in four lineages of *S. polyclona*

Even among closely related lineages, differences in the evolutionary patterns of SLRs are observed in E, E-TS, W1, and W2. To understand origin and evolutionary history of the current sex chromosome of *S. polyclona*, we observed the clustering of gametologs within SLRs of *S. polyclona* complex and their relative species. A total of 62 phylogenetic trees were obtained using the sex-linked single-copy orthogroups in *S. polyclona*, *S. purpurea*, and *S. arbutifolia*. Trees of the two ancient sex-linked single-copy orthogroups (fully sex-linked genes shared by 15XY and 15ZW species of *Vetrix*-clade ^12^) revealed that the 15W-linked gene of W2, E-TS, *S. purpurea*, and the 15X-linked gene of *S. arbutifolia* cluster together, while the 15Z-linked gene of W2 (Z(b), Z(c), Z(d)), E-TS, *S. purpurea*, the 15Y-linked gene of *S. arbutifolia*, and the 15Z and 15W-linked genes of W1 and E form a well-supported separate clade (Fig. 3(a), Extended Data Fig. 1(a)). We also obtained four other trees that 15X and 15Y-linked genes of *S. arbutifolia* formed a clade sister to 15Z and 15W-linked genes of the complex and *S. purpurea* (Fig. 3(b), Extended Data Fig. 1(b)∼(d)). Within the complex-*S. purpurea* subclade, 15W-linked genes of *S. purpurea*, W2, and E-TS cluster together, while 15Z-linked genes of *S. purpurea*, W2 (Z(b), Z(c), Z(d)) and E-TS, and 15Z and 15W-linked genes of W1 and E formed a well-supported clade. The results support that the 15ZW species of the *Vetrix* clade arose from an 15XY ancestor ^12^, and ancestral 15ZW species evolved fully sex-linked genes before the 15ZW species split. These results also suggest that occasional recombination occurred between 15Z and 15W of W1 and E, or E and W1’s 15W arose from 15Z. For the other 56 sex-linked single-copy orthogroups, the distributions of X, Y, Z, and W gametologs are disordered. We did not find any sex-linked single-copy orthogroup tree that support 15Z and 15W-linked genes of W1 and E forming a clade with W-linked genes of W2 and E-TS. Furthermore, we observed similar distribution pattern of male specific sequences (partial *ARR17*-like duplicates) in both 15Z-SLR and 15W-SLR of lineages E and W1, and the phylogenetic tree of intact *ARR17*-like genes is consistent with these six trees constructed with ancient sex-linked genes (Fig. 3(c), Fig. 4, Supplementary Fig. 11, see below). These results further imply that either 15W of E and W1 arose from 15Z, or they experienced occasional recombination.

**Figure 3.**
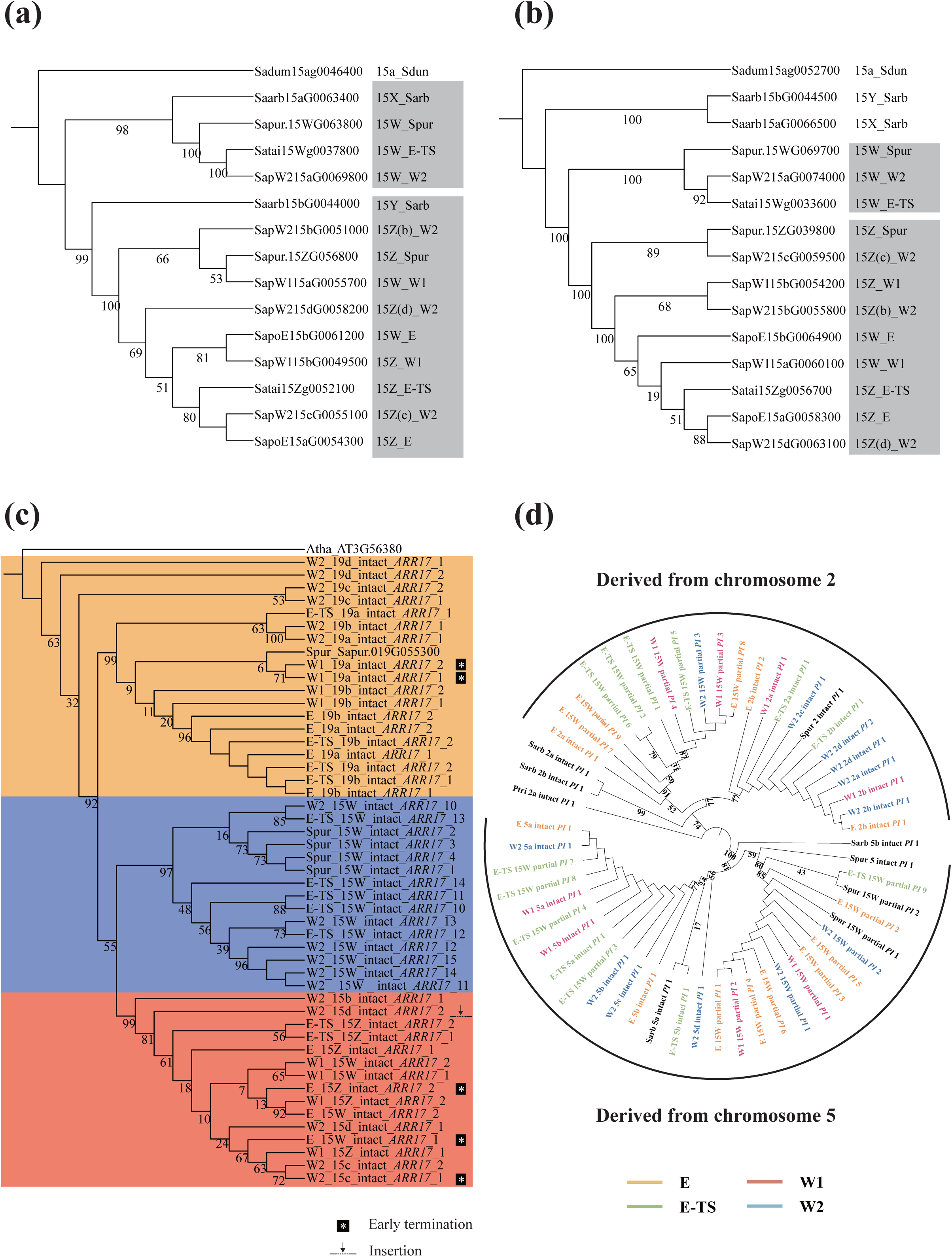
The phylogeny of ancient sex-linked genes, intact *ARR17*-like duplicates and *PI*-like duplicates. (a) Ancient sex-linked genes in Topology 1 derived before the speciation of *S. arbutifolia*. For the other ancient sex-linked genes in Topology 1, see Extended Data Fig. 1a. (b) The ancient sex-linked genes in Topology 2 originated after the speciation of *S. arbutifolia*, but before the speciation of *S. polyclona* complex. For the other three ancient sex-linked genes in Topology 2, see Extended Data Fig. 1b,1c, and 1d. (c) The phylogenetic tree of intact *ARR17*-like genes in E, E-TS, W1, and W2. The orange and purple background color shows that the *ARR17*-like genes on chromosome 19 and 15W-SLR. The red background color shows that the *ARR17*-like genes on 15Z-SLR (turnovers in E and W1), which duplicated and translocated from 15W-SLR. The early termination and insertion were marked after the *ARR17*-like genes. (d) The phylogenetic tree of intact and partial *PI*-like duplicates in the *S. polyclona* complex, *S. purpurea*, *S. arbutifolia*, and *Populus trichocarpa*. Numbers marked on the tree represent bootstrap values. E, E-TS, W1, W2: *S. polyclona*-E, *S. polyclona*-E-TS, *S. polyclona*-W1, *S. polyclona*-W2; Sarb: *S. arbutifolia*; Spur: *S. purpurea*; Sdun: *S. dunnii*; Ptri: *P. trichocarpa*.

**Figure 4.**
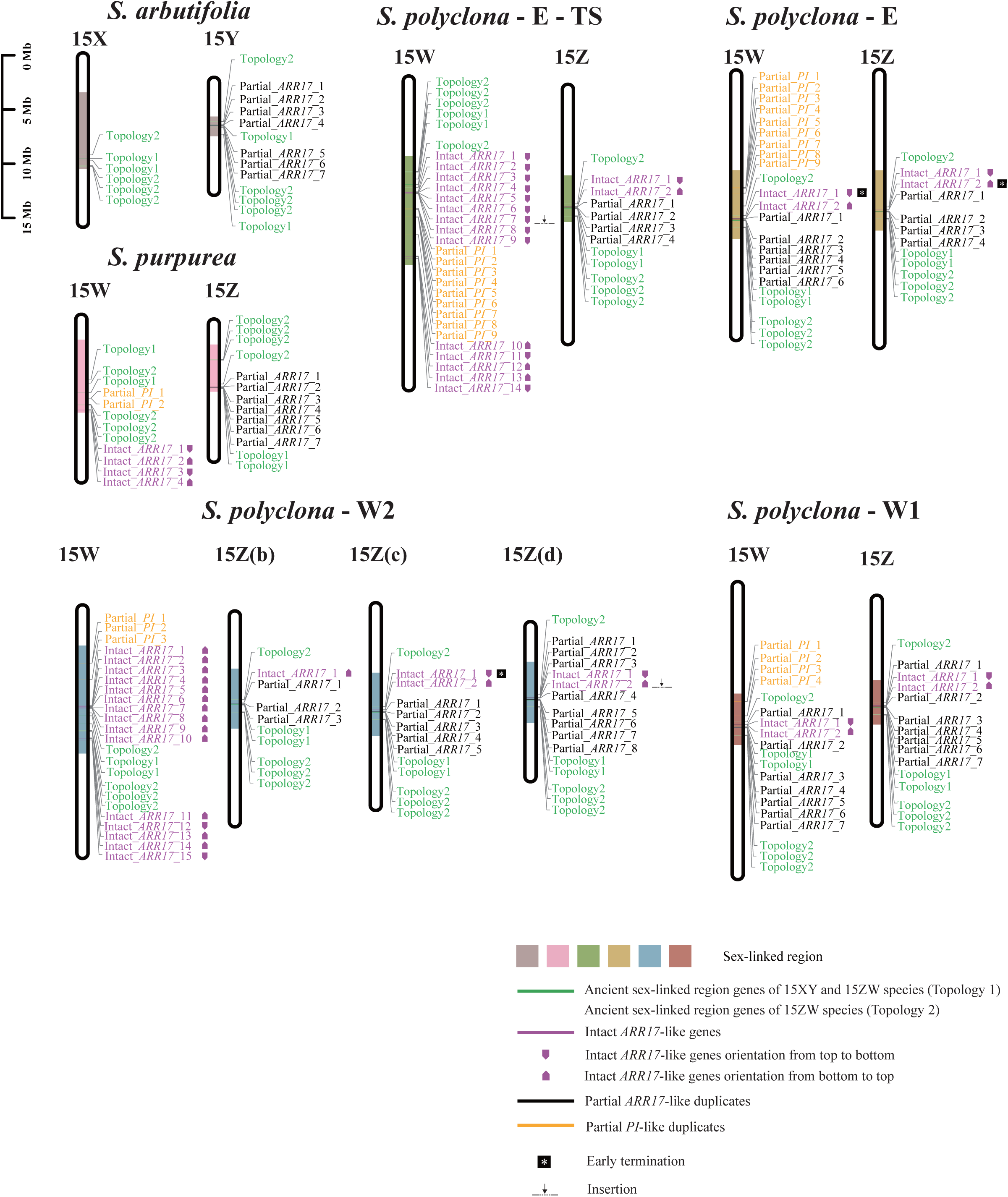
The distribution of sex-linked genes and candidate sex-determination factors on sex chromosomes in willows. The colored regions on chromosomes are SLRs. Green markers represent the ancient sex-linked genes, Topology 1 and Topology 2 in Fig. 3(a)(b) and Extended Data Fig. 1. Magenta, black, and yellow markers show the intact *ARR17*-like duplicates, partial *ARR17*-like duplicates, and partial *PI*-like duplicates, respectively. Early termination and insertion were marked after the *ARR17*-like genes. The magenta arrows indicate the direction of intact *ARR17*-like duplicates (the tip is the end of the gene).

We identified *ARR17*-like duplicates in *S. polyclona* complex, which is involved in sex determination of *Salix* and *Populus* ^10,13^. The intact *ARR17*-like genes are distributed on chromosome 19 of *S. chaenomeloides*, *S. dunnii*, *S. arbutifolia*, *S. purpurea*, *S. babylonica*, and on 15W-SLR of *S. purpurea* ^12,13,15,38^. Consistent with this, we observed intact *ARR17*-like genes on chromosome 19 and 15W-SLRs in the *S. polyclona* complex. However, we also detected intact *ARR17*-like genes on 15Z-SLRs in the complex (Extended Data Table 5), which has this far not been observed in any other Salicaceae species. The intact *PI*-like duplicates, as the downstream targets of intact *ARR17*-like duplicates, were found on autosomes 2 and 5 of *S. polyclona* complex, similar to *S. babylonica* ^15^. In fact, autosomes 2 and 5 in *Salix* are homologous chromosomes that originated from a whole genome duplication event, as reported in previous studies ^15,39^. However, additional partial *PI*-like duplicates were distinguished on 15W-SLRs on *S. purpurea* and *S. polyclona* complex, whereas there were no partial *PI*-like duplicates on SLRs of *S. babylonica* and *S. arbutifolia* (Extended Data Table 5).

In addition, E, E-TS, W1, and W2 have accumulated different female and male specific genes, respectively (Supplementary Fig. 12, Extended Data Table 6), suggesting their SLRs experienced diverse evolutionary pressures. Based on the annotation results, we identified several candidate genes associated with flowering. The *VAL1*- and *VAL2*-like genes ^40^ (SapW115aG0051300 and SapW115aG0051400) in 15W-SLR of W1 could indirectly accelerates flowering by influencing the expression of *FLC*. We also identified *HYL1*-like gene (SapW115bG0036300) in 15Z-SLR of W1, which is related to stamen structure ^41^. In addition, the *RBR*-like (SapW215aG0049700) and *ARF3*-like (SapW215aG0051700) genes were found in 15W-SLR of W2, which promote flower organogenesis ^42,43^. Furthermore, the four lineages accumulated different inversions in their SLRs (Fig. 2(b)).

We also found that W2 has experienced rediploidization. The collinearity between 15Z(b)-SLR and 15Z(c)-SLR is similar in W2, while different with 15Z(d)-SLR (Supplementary Fig. 10(g)). The number of partial *ARR17*-like duplicates on 15Z(b)-SLR (3) and 15Z(c)-SLR (5) is less than on 15Z(d)-SLR (8) (Extended Data Table 5). In addition, the average haplotype genome size and the number of mRNA of W2 decreased compared to other diploid lineages (Supplementary Table 9).

### The origin and evolutionary trajectory of *ARR17*-like and *PI*-like duplicates

To clarify the origin and evolutionary trajectory of sex-determining genes, we reconstructed the phylogenetic tree of intact *ARR17*-like and *PI*-like duplicates in the *S. polyclona* complex, and found extensive translocations of intact *ARR17*-like genes (Fig. 3(c)(d), Supplementary Fig. 11). As in other willow species, intact *ARR17*-like genes are likely ancestral on chromosome 19 in *S. polyclona*, and translocated from there to 15W-SLR. Subsequently, the intact *ARR17*-like genes experienced duplication and translocation again, copying from 15W-SLR to 15Z-SLR. Like ancestral sex-linked genes (Fig. 3(a)(b), Extended Data Fig. 1), the intact *ARR17*-like genes on 15W-SLR and 15Z-SLR of E and W1 cluster with other intact *ARR17*-like genes on 15Z-SLR of E-TS and W2 (Fig. 3(c)), suggesting either that recombination occurs on E and W1 or neo-W chromosomes evolved in them. In addition, the number of intact *ARR17*-like genes on the 15W-SLR of E-TS (14 copies) and W2 (15 copies) has greatly expanded. These *ARR17*-like genes have two origins, with some duplicated and translocated from chromosome 19, and the others derived from the duplication and translocation within 15W-SLR (Intact_*ARR17*_1-9 in E-TS and W2) (Supplementary Fig. 11, Extended Data Table 5).

Some of the partial *PI*-like duplicates on 15W-SLRs of the *S. polyclona* complex clustered with intact *PI*-like duplicates on autosomes 2 of *S. polyclona* complex, *S. purpurea*, and *S. arbutifolia*, and the other clustered with intact *PI*-like duplicates on autosomes 5 of *S. polyclona* complex, *S. purpurea*, and *S. arbutifolia*, suggesting that the 15W-linked partial *PI*-like duplicates originated from autosomes 2 and 5 (Fig. 3(d)).

### Sex determination in the *S. polyclona* complex

All of the possible sex-determining factors including ancient sex-linked genes, intact *ARR17*-like duplicates, partial *ARR17*-like duplicates, partial *PI*-like duplicates were identified on the sex chromosomes of *S. polyclona* complex, *S. arbutifolia* and *S. purpurea* (Fig. 4). We explored the sex determination strategy of *S. polyclona* complex according to the distribution and expression of candidate sex-determining genes. Surprisingly, the ancestral Salicaceae sex determination factor intact *ARR17*-like duplicates were barely expressed in the flower buds and catkins of *S. polyclona* complex (Fig. 5). Some intact *ARR17*-like duplicates exhibit early termination or insertion (Fig. 3(c), Fig. 4, Supplementary Fig. 11, Supplementary Fig. 13), which may affect their function, but the other intact *ARR17*-like duplicates remain unexpressed, and several others have very low expression levels. Given this expression pattern, it is likely that the *ARR17*-like genes no longer act as a key sex determination factor in *S. polyclona*.

**Figure 5.**
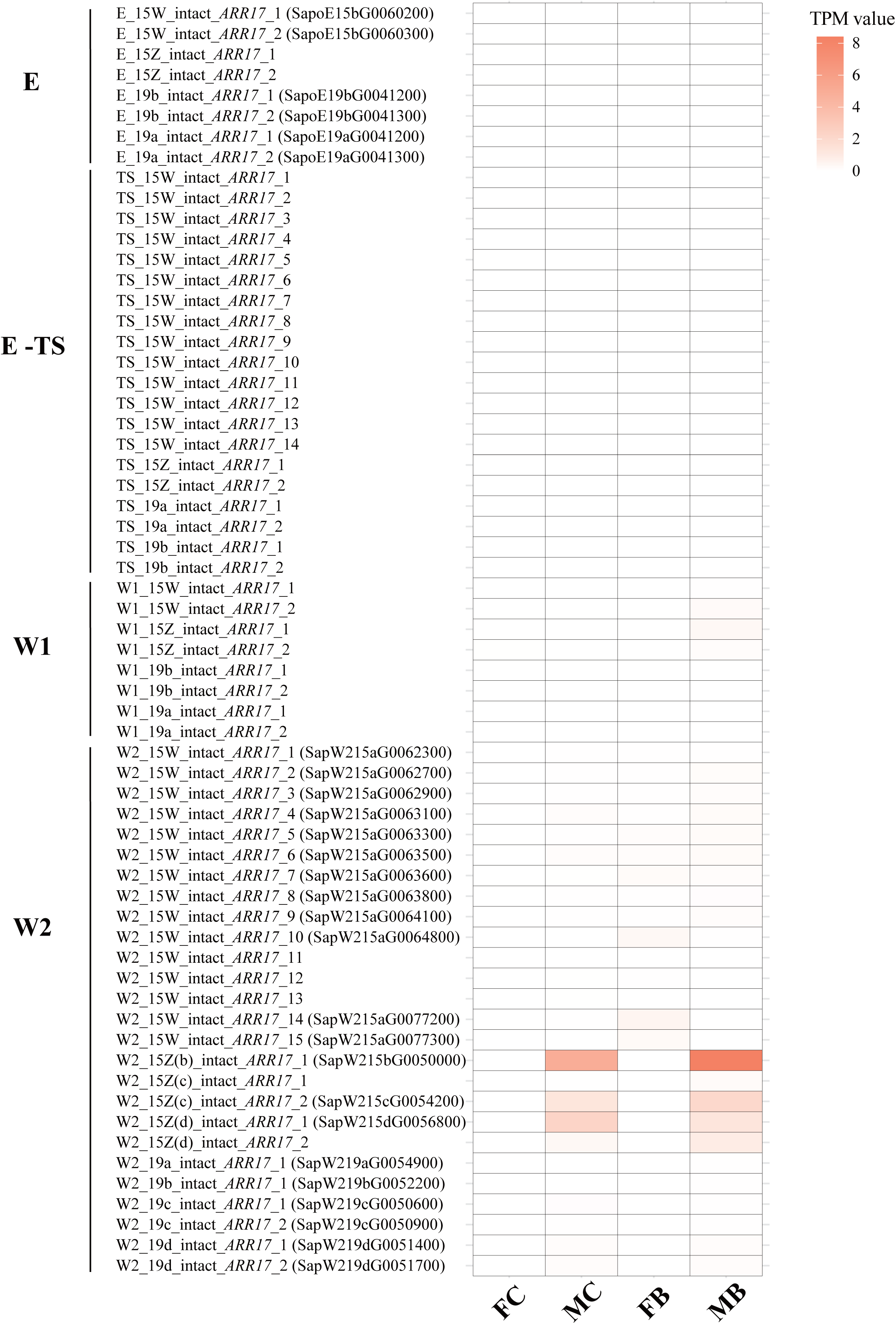
The expression (TPM) of intact *ARR17*-like duplicates of *S. polyclona* complex. No annotation results are available for duplicates without a given gene ID. E, E-TS, W1, W2: *S. polyclona*-E, *S. polyclona*-E-TS, *S. polyclona*-W1, *S. polyclona*-W2.

The *ARR17*-like genes determine sex indirectly by hindering the male factor B-class gene *PISTILLATA* (*PI*), which in turn determines femaleness ^19^. In the *S. polyclona,* we observed high expression of intact *PI* duplicates in male buds and catkins, but hardly any in female buds and catkins (Fig. 6(a)), suggesting *PI* still plays a key role in sex determination in the complex. It is worth noting that additional partial *PI*-like duplicates distributed on 15W-SLRs of *S. polyclona* complex could play a crucial role in the sex determination pathway (Fig. 4, Extended Data Table 5). Moreover, we found a large accumulation of sRNAs near the partial *PI*-like duplicates of 15W-SLRs of the four lineages (Fig. 6(b)). This expression pattern indicates that the partial *PI*-like duplicates likely inhibit the expression of intact *PI*-like duplicates by producing small RNAs, and thus have emerged as a new female factor in the complex.

**Figure 6.**
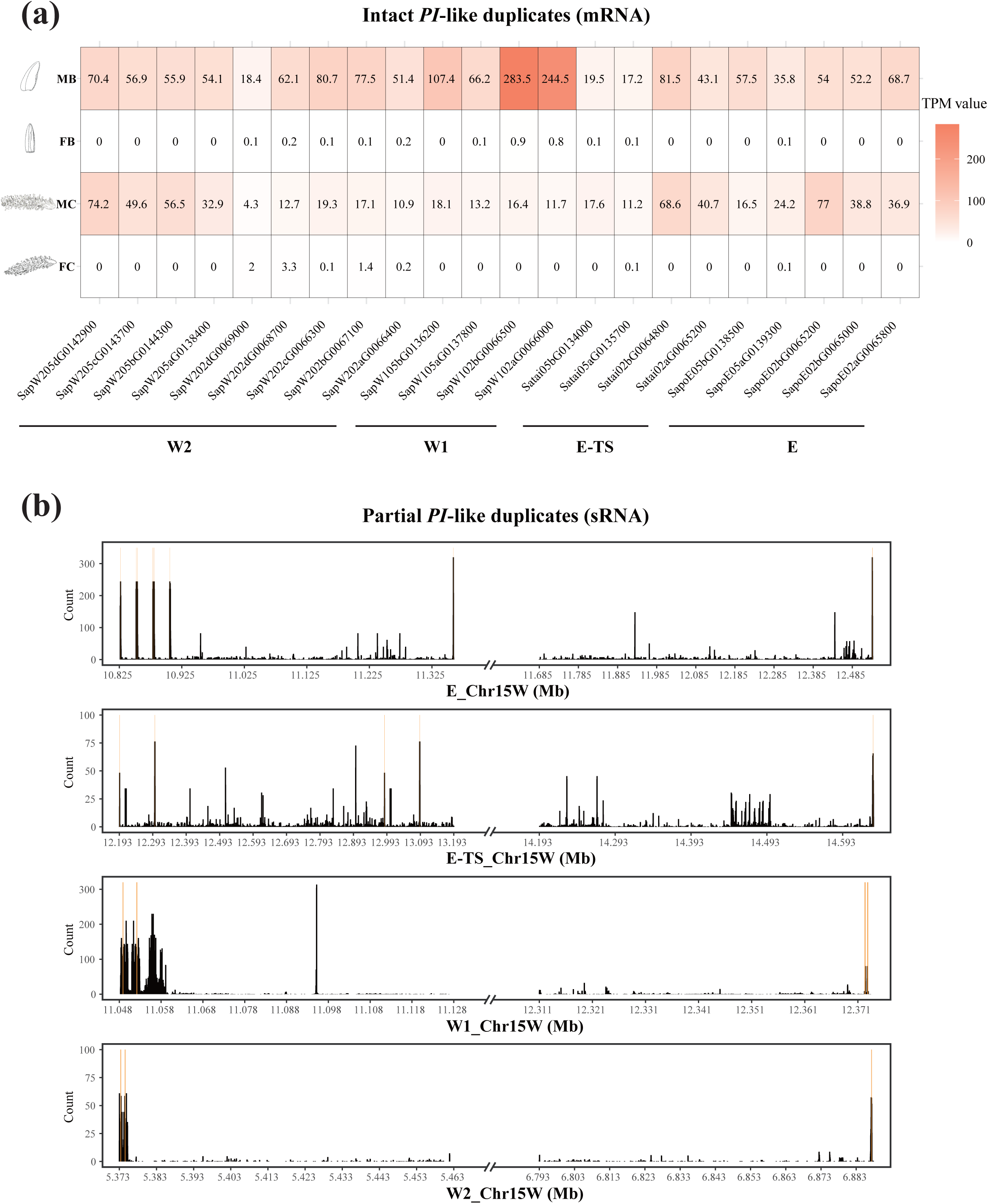
The expression pattern of *PI*-like duplicates in the *S. polyclona* complex. (a) The mRNA expression (TPM value) of intact *PI*-like duplicates in male buds (MB), female buds (FB), male catkins (MC), and female catkins (FC). (b) The distribution of small RNAs around the partial *PI*-like duplicates. The yellow lines marked the position of partial *PI*-like duplicates.

Finally, we inferred the sex determination model of *S. polyclona* complex (Fig. 7). Although there are various duplications and translocations for upstream sex determination factor intact *ARR17*-like duplicates, they may be taken over by the partial *PI*-like duplicates on 15W-SLRs, which were recruited from autosomes 2 and 5. For female individuals (ZW or ZZZW), the partial *PI*-like duplicates on 15W-SLRs can produce sRNA to inhibit the intact *PI*-like duplicates on autosomes 2 and 5. In contrast, the intact *PI*-like duplicates can function normally in male individuals (ZZ or ZZZZ).

**Figure 7.**
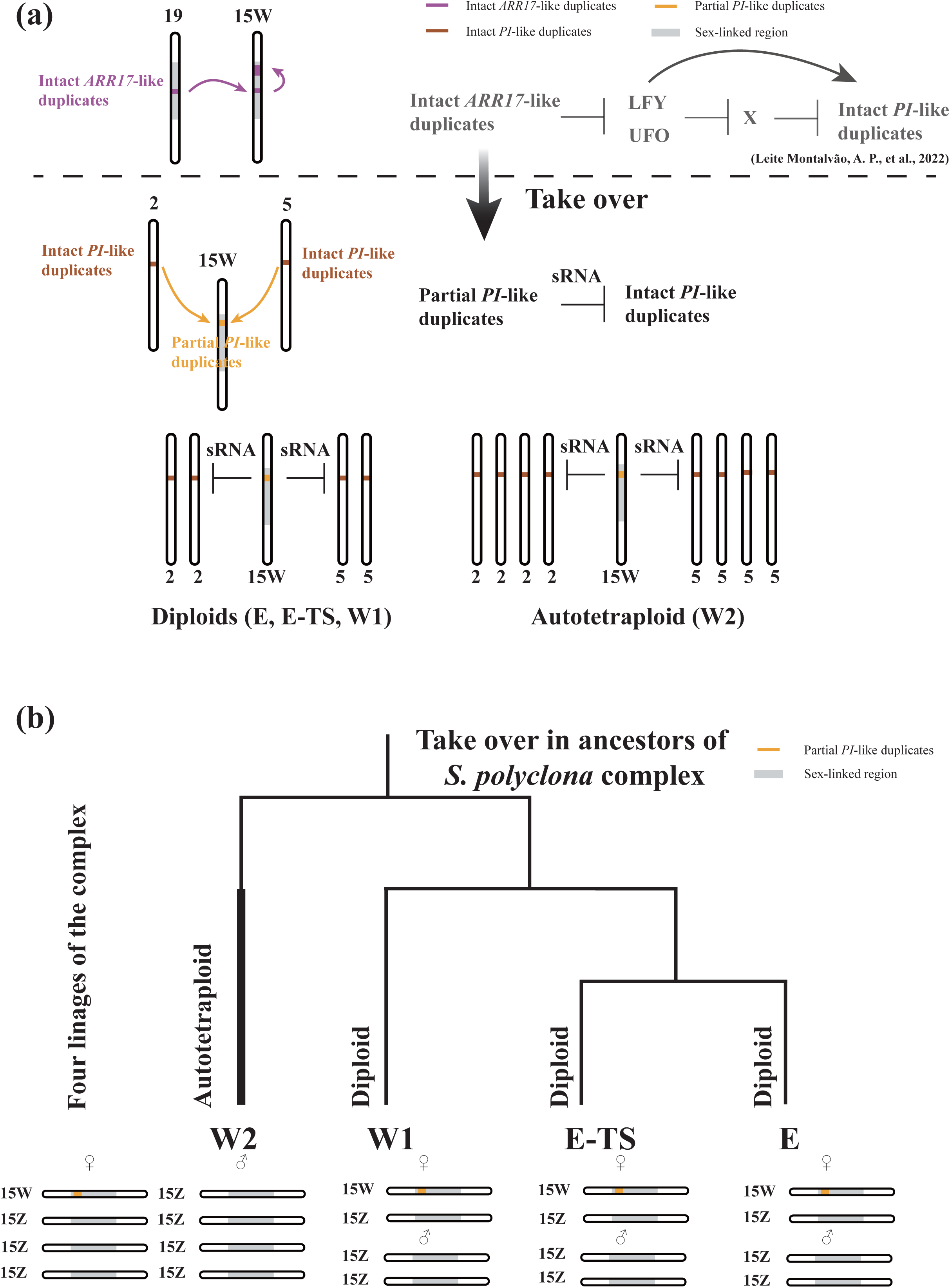
The sex determination model in the *S. polyclona* complex. (a) The evolutionary trajectories of sex determination factors and the potential sex determination model in the *S. polyclona* complex. The upstream apical sex-determining factor intact *ARR17*-like duplicates were replaced by partial *PI*-like duplicates. Magenta and yellow markers show the intact *ARR17*-like duplicates and partial *PI*-like duplicates, respectively. (b) The simplified phylogenetic tree of the *S. polyclona* complex. The partial *PI*-like duplicates take over from the original apical sex-determining gene.

This is a newly sex determination strategy in Salicaceae, which is different from the previous turnovers mediated by apical translocation of sex determination gene ^10–17^. The sex determination pathway of the *S. polyclona* complex has diverged from its ancestors, with negative regulation of downstream genes (partial *PI*-like duplicates) now taking over the core upstream gene (intact *ARR17*-like duplicates).

## Discussion

In this study, we assembled four haplotype-resolved genomes of the *S. polyclona* complex, including three diploids (E, E-TS, and W1) and an autotetraploid (W2), and found that frequent translocations of intact *ARR17*-like duplicates occurred in all of them. More importantly, a new putative sex-determining gene, partial *PI*-like duplicates, was detected in the *S. polyclona* complex, which replaced the function of intact *ARR17*-like duplicates (Fig. 7). This study provides a novel example about the recruitment and replacement of new sex-determining loci, and suggests a downward trajectory of the sex-determination pathway, different from the upward trajectory suggested by Wilkins^22^.

The ancestral sex-determining mechanism in poplars and willows is that partial *ARR17-* like gene duplicates produce small RNAs silencing intact *ARR17*-like genes, indirectly activating intact *PI*-like genes and thereby determining males ^10–17,19^ In contrast, expressed intact *ARR17*-like genes indirectly suppress intact *PI*-like genes and determine females ^10,15,19^. Despite broad conservation of this mechanism in other poplars and willows, we find that intact *ARR17*-like duplicates are barely expressed in females of the *S. polyclona* complex (Fig. 5). Despite this, downstream intact *PI*-like duplicates in the females of *S. polyclona* complex are barely expressed, consistent with the expression pattern of *S. babylonica* females ^15^ (Fig. 6(a)). Furthermore, we detected large accumulation of sRNAs near the partial *PI*-like duplicates of 15W-SLRs of the four lineages of the complex. This suggests that the ancestors of *S. polyclona* complex recruited the new sex-determining factor partial *PI*-like duplicates from autosomes 2 and 5, which could replace the function of *ARR17*-like duplicates, inhibiting downstream intact *PI*-like duplicates.

We also found two partial *PI*-like duplicates in 15W-SLR of *S. purpurea* (Fig. 4), however Hyden proposed that *ARR17* and *GATA15* determine the sex of *S. purpurea* ^44^. Hence, it is unknown whether the newly found sex determination mechanism of the *S. polyclona* complex is a common origination in *Vetrix*-clade 15ZW species, so this needs to be tested with more genomic data in the future.

A previous study proposed that the translocation and duplication of intact *ARR17*-like genes induce the sex chromosome turnover in *S. purpurea* (15XY→15ZW) ^13^, while we detected a new turnover model in the *S. polyclona* complex. The new sex-determining factor, partial *PI*-like duplicates, appears to replace the function of original apical sex-determining gene of Salicaceae, *ARR17*-like duplicates (Fig. 7). This change involved a shift from intact *ARR17*-like duplicates on 15ZW SDS to partial *PI*-like duplicates on 15ZW SDS.

The partial *PI*-like duplicates on 15W-SLR are analogous to *DMW* in *Xenopus*^45^, partial copies of *DMRT1* that actually inhibit *DMRT1* to determine female function. Similarly, partial *PI-*like duplicates act as negative regulators of downstream sex determining gene (intact *PI*-like duplicates). More importantly, our results suggest that sex chromosome turnover in plants can be driven by rewiring of the sex determining pathway in similar ways to that observed in animals. It remains to be determined whether sex chromosome turnovers are more often driven by translocations or duplications of existing apical sex-determining genes ^6,8^ or by rewiring of the pathway itself.

## Materials and Methods

### Plant material

We collected leaves from a female plant of each four lineages, W2 (HL00183), W1 (HL00215), E (HL00164), and E-TS (YGDF2) of *Salix polyclona* diploid-autotetraploid complex for whole genome sequencing ^35^. Catkin, flower bud, young leaf, and stem samples from the four female plants and male bud and catkin samples from each of the four lineages were collected for RNA sequencing. Samples were frozen in liquid nitrogen and stored at −80°C until total DNA or RNA extraction. Leaves of ten males of E-TS, three females and 14 males of E, 14 males of W1, and nine males of W2 were sampled and dried in silica-gel for whole genome resequencing and ploidy estimation. Voucher specimens of these new sequenced samples are deposited in the herbarium of Shanghai Chenshan Botanical Garden (CSH). We also downloaded genome sequence data of four E-TS, 22 W1, 19 E, nine W2 individuals of the complex published by ^35^. Detailed information of these datasets was listed in Extended Data Table 7.

### Ploidy determination

The ploidy levels of 31 individuals of the *S. polyclona* complex were measured by flow cytometry, using diploid *S. baileyi* (2x = 2n = 38 ^46^) as an external standard. We followed the protocol of ^47^. We chopped leaf tissue with a razor blade after incubated it for 80 min in 1 mL LB01 buffer. A 38-μm nylon mesh was then used to filter the homogenate. The filtered homogenate was treated with 80 μg/mL propidium iodide (PI) and 80 μg/mL RNase to stain the nuclei. Estimations were done in MoFlo-XDP flow cytometer using Summit v.5.2 (Beckman Coulter Inc.). The ploidy level was calculated as sample ploidy = reference ploidy × mean position of the sample peak/mean position of reference peak.

We used JELLYFISH ^48^ to construct *k-mer* frequency distributions of tetraploid *S. polyclona*-W2 (HL00183) (*k-mer* = 21). Genomescope 2.0 ^49^ was used to distinguish autotetraploidy from allotetraploidy according to the patterns of nucleotide heterozygosity. Due to preferential pairing between bivalents formed by the same subgenome in allotetraploids, the allotetraploids should have a higher proportion of aabb than aaab (aabb > aaab), while autotetraploids should have a higher proportion of aaab (aaab > aabb) because of quadrivalents during meiosis ^49^.

### Genome sequencing

We sequenced 50 individuals (41 diploids and 9 tetraploids) of the *S. polyclona* complex. Genomic DNA from these 50 samples was extracted from leaves using the Magnetic Plant Genomic DNA Kit (Tiangen, China). Paired-end libraries were constructed for all samples. Whole-genome sequencing with an expected depth of 20× for diploids and 80× for tetraploids were performed on Illumina NovaSeq 6000 and NovaSeq X Plus by Beijing Novogene Bioinformatics Technology.

For long-read sequencing, PacBio HiFi libraries were prepared from genomic DNA. We extracted genomic DNA from four lineages of the *S. polyclona* complex using CTAB method, and the primary band was larger than 30 kb. Then, PacBio large insert libraries were created by SMRTbell Express Template Prep Kit 2.0. These libraries were sequenced by Novogene using the PacBio Sequel II platform with the Circular Consensus Sequencing (CCS) model. In order to improve the accuracy of the assembly, we also performed ONT sequencing for autotetraploid *S. polyclona*-W2, and DNA was extracted using phenol–chloroform. ONT libraries were prepared following the Nanopore 1D Genomic DNA by ligation protocol, and sequenced by Novogene on the PromethION platform.

Hi-C libraries were constructed using standard procedures ^50^. Tender leaves from four lineages of the *S. polyclona* complex were used for library preparation. The leaves were fixed with a 4% formaldehyde solution, and then the cross-linked DNA was isolated from nuclei. Subsequently, DNA was digested with restriction enzyme MboI, and restriction fragment ends were biotinylated, purified and ligated. Hi-C libraries were controlled for quality and sequenced on NovaSeq 6000 by Novogene.

### RNA sequencing

Total mRNA was extracted from catkin, flower bud, young leaf, and stem samples from the four female plants and male catkin samples from each four lineages using a Polysaccharide Polyphenol Plant Total RNA Extraction Kit (Tiangen, China). The integrity of RNA was assessed using the Fragment Analyzer 5400 (Agilent Technologies, CA, USA), and then libraries were generated using NEBNext® UltraTM RNA Library Prep Kit for Illumina® (NEB, USA).

We also performed small RNA sequencing for female and male flower buds from our four lineages. Total RNA was extracted using RNAprep Pure Plant Plus Kit (Tiangen, China). After adaptors were ligated to the ends of the small RNA molecules, reverse transcription primer hybridization was used to create the first strand cDNA. The double-stranded cDNA library was created using PCR enrichment. Libraries with insertions ranging from 18 to 40 bp were prepared for sequencing following size selection and purification. The mRNA-seq and small RNA-seq sequencing were performed on an Illumina NovaSeq 6000 by Novogene.

### Genome assembly

We used a similar strategy to assemble and annotate the genome assemblies of the four lineages of *S. polyclona* complex (E, E-TS, W1, W2). Initial contigs were assembled based on PacBio HiFi reads using hifiasm ^51^ for *S. polyclona*-E, *S. polyclona*-E-TS and *S. polyclona*-W1. For autopolyploid *S. polyclona*-W2, ONT reads were also used to assemble initial contigs. Then Hi-C reads were aligned to the haplotype contig genome using Juicer ^52^ in order to facilitate chromosome-level genome assembly. Then, using 3d-dna ^53^, a preliminary Hi-C-assisted chromosomal assembly was completed. Subsequently, manual inspection and correction were conducted using Juicebox ^54^. The primary objectives of this process were to refine chromosomal boundaries, eliminate improper insertions, modify orientations, and fix assembly faults. Using the LR_Gapcloser ^55^, gap filling based on HiFi readings was carried out in order to improve the assembly.

GetOrganelle ^56^ was used to assemble chloroplast and mitochondrial genomes. The fragmented contigs were aligned to the chromosome-level genome and organelle genome sequences by Redundans ^57^, identifying redundant segments. Moreover, rDNA fragments and low-coverage fragments or haplotigs among the dispersed sequences were eliminated. Furthermore, our short-read data was used to polish with Nextpolish ^58^ for genome base correction.

In order to improve our chromosome-scale, haplotype-resolved genome assemblies of *S. polyclona*-E, *S. polyclona*-E-TS, *S. polyclona*-W1, *S. polyclona*-W2, the reads mapped near the telomeres were selected and assembled into contigs using hifiasm, and then these contigs were mapped back to the chromosomes. The chromosomes were compiled as chromosome 01 to chromosome 19 according to the homologous relationship with *S. brachista* ^59^. We obtained two haplotypes (a, b) for *S. polyclona*-E, *S. polyclona*-E-TS, *S. polyclona*-W1, and four haplotypes (a, b, c, d) for *S. polyclona*-W2. There is an extra chromosome 20 in *S. polyclona*-W2.

### Genome annotation

Coding gene prediction was accomplished by combing evidence from homology-based prediction, transcript prediction, and *de novo* prediction strategies. For homology-based prediction, we used the publicly available Salicaceae protein sequences as homologous protein evidence for gene annotation. For transcript prediction, we assemble transcripts using Trinity ^60^ and StringTie ^61^, and removed redundancy with CD-HIT ^62^ (identity > 95%, coverage > 95%). PASA pipeline ^63^ was used to annotate gene structure based on transcriptome data, and full-length genes were identified by aligning with homologous protein. These full-length genes were used for AUGUSTUS ^64^ training, and conducted five replicates of optimization. Then gene structures were predicted by MAKER ^65^ pipeline based on the repeat-masked genome with *de novo* prediction, transcript and homolog protein evidence. Next, MAKER and PASA annotation results were integrated to produce consensus gene sets using EVidenceModeler (EVM) gene structure annotation tool ^66^. Finally, untranslated regions (UTRs) and alternative splicing were annotated using PASA ^63^, and genes with less than 50 amino acids and missense annotations (internal stop codon or ambiguous base, no start codon or stop codon) were removed.

Non-coding RNAs (ncRNAs) were annotated using tRNAScan-SE ^67^, RfamScan, and Barrnap(https://github.com/tseemann/barrnap). Gene functions were annotated based to homology and similarity searches. In addition to the eggNOG-mapper ^68^ annotation, DIAMOND ^69^ was used to sequence similarity searches (identity > 30%, E-value < 1e-5), and protein databases including Swiss_Prot, TrEMBL and NR. InterProScan ^70^ was used to domain similarity search. Repeat elements were identified by EDTA ^71^ (--sensitive 1 --anno 1). Then, RepeatMasker (http://www.repeatmasker.org/RepeatMasker/) was used to determine repetitive regions within our genome assemblies. BUSCO was used to assess genome completeness (embryophyta_odb9 database).

### Phylogenetic analysis

We performed a phylogenetic analysis of the eight willow genomes including *S. dunnii* (haplotype *a*), *S. arbutifolia* (haplotype *a*), *S. purpurea*, *S. brachista*, *S. polyclona*-W2 (haplotype *a*), *S. polyclona*-W1 (haplotype *a*), *S. polyclona*-E-TS (haplotype *a*), *S. polyclona*-E (haplotype *a*), and used *Populus trichocarpa* as outgroup (Supplementary Table 4). Single-copy homologous protein sequences were identified using OrthoFinder ^72^. In order to avoid the effects of sex chromosomes in other *Salix* and *Populus* species on the tree, the sequences on chromosomes 7, 15 and 19 were removed in this analysis. Then, these protein sequences were aligned using MAFFT ^73^, and we reconstructed gene trees with IQ-TREE (-m MFP -bb 1000 -bnni -redo) ^74^. The species tree was estimated with ASTRAL ^75^ according gene trees. The species tree was then used as an input tree to estimate the divergence time using MCMCTREE in PAML ^76^ package (burnin = 400000, sampfreq = 10, nsample = 100000). We used two fossils and one inferred time at three nodes: (1) the root node of Salicaceae (48 Mya), (2) the divergence time between *Salix* and *Vetrix* (37.15 Mya to 48.42 Mya), and (3) the ingroup of *Vetrix* (*Chamaetia*-*Vetrix*) clade (23 Mya) ^77^.

### Reads mapping and variant calling

Fastp ^78^ was used to filter all sequence reads, and clean reads were used for subsequent analysis. The BWA-MEM algorithm from bwa 0.7.12 ^79,80^ was used to align clean reads to each genome (both haplotypes for diploid W1 and E, and the *a* & *b* and *c* & *d* haplotypes of W2) with default parameters. Samtools 0.1.19 ^81^ was used to deal with the mapped data. PCR replicates were filtered using sambamba 0.7.1 ^82^. GATK 4.2.2.0 version was used to call variants. Hard filtering was carried out with parameters “QD<2.0, FS>60.0, MQ<40.0, MQRankSum<–12.5, ReadPosRankSum< –8.0, and SOR>3.0”. Only biallelic sites were used in subsequently filtering steps. Sites with coverage above approximately twice the mean depth at variant sites across all samples were discarded. Sites with sequencing depth < 4× were treated as missing, and sites with missing samples > 10% or with minor allele frequency <0.05 were filtered.

### Identification of the SLRs of the four lineages of *S. polyclona* complex

We identified the SLRs of each sample of the *S. polyclona* complex using a combination of the results of CQ, *F*_ST_, the location of inversions, and syntenic relationships between the identified regions. The CQ method ^37^ was used to detect Z and/or W sex chromosomes of the four lineages using cq-calculate.pl software based on clean reads. We calculated the CQ for each 50-kb nonoverlapping window of the genomes based on combined female and male clean read data sets of the four lineages, respectively. For female heterogamety, the CQ is the normalized ratio of male to female alignments, and CQ value close to 2 (for ZW/ZZ) in windows in Z-linked region and to zero in windows in the W-linked region. The Changepoint package ^83^ was used to detect the boundaries of the SLRs based on CQ values between the sexes of *S. polyclona* complex.

Weighted *F*_ST_ values between the sexes of E, W1 and W2 were calculated using the ^84^ estimator with 100 kb windows and 10 kb steps, respectively. We used the changepoint package ^83^ to assess significance of differences in the mean and variance of the *F*_ST_ values between the sexes of chromosome 15 windows, which was revealed as sex chromosome of the complex ^35^. The E-TS lineage includes a very small population on the top of Tai Mountain, and only 34 individuals were recorded before ^85^. We just obtained re-sequencing datasets from four female and 10 males individuals ^35^, hence we didn’t calculate the *F*_ST_ of them.

We also conducted synteny analysis between the sex chromosomes of each lineage of the complex using Python version of MCScan ^86^ with parameter “--cscore=.99”. The boundaries of inversions identified by synteny were contained within SLRs. The corresponding SLRs between sex chromosomes 15Z and 15W were detected according to the synteny results.

### Ancient sex-linked genes identification of the complex

For species that shared common ancestral SLRs, their ancestral X and Y or W and Z alleles are expected to cluster by gametologs ^12,87^. The species with a 15ZW system of the *Vetrix*-clade were suggested to have arisen from a common 15XY ancestor. We expect that there are more genes clustering by gametologs within 15ZW species compared to the clustering of genes between 15ZW species and 15XY relatives.

The OrthoFinder ^72^ was employed to identify single-copy genes in SLRs of 15W and 15Z of *S. polyclona* complex, *S. purpurea*, 15X and 15Y of *S. arbutifolia*, and chromosome 15a of *S. dunnii*. We used MAFFT ^73^ to align the obtained single-copy homologous genes, and IQ-TREE ^74^ to reconstruct phylogenetic trees using *S. dunnii* as outgroup.

### *ARR17*-like and *PI*-like duplicates identification and phylogeny

Intact *ARR17*-like genes (including intact 5 exons) and partial *ARR17*-like duplicates (including less than 5 exons) were identified using the same strategy as in other willows ^12,13,15^. We used BLASTN to blast *ARR17*-like gene (Potri.019G133600 ^10^, including five exons) in the *S. polyclona* complex and *S. purpurea* (-evalue 1e-5 -word_size 8). To ensure that all the regions of *ARR17*-like duplicates were identified, we extracted the aligned regions and 200 bp upstream and downstream of these regions. Then, Geneious Prime 2023.2.1 (https://www.geneious.com/) was used to align and visualize sequences.

The five exons of *ARR17*-like genes were identified and annotated using the intact *ARR17*-like gene in *S. purpurea* (Sapur.019G0055300), then all the *ARR17*-like duplicates were aligned from the start codon ATG. The coding sequence (CDS) of intact *ARR17* genes were used to construct a phylogenetic tree with IQ-TREE ^74^, using the *ARR17* gene in *Arabidopsis thaliana* (AT3G56380) as outgroup. Because a large number of tandem duplications can affect the topology of the phylogenetic tree ^88^, we independently reconstructed phylogenetic tree for the intact *ARR17* genes on 15W-SLRs of E-TS and W2. We also used the same method to identify the *PI*-like genes (7 exons) in the *S. polyclona* complex, *S. purpurea*, and *S. arbutifolia* genomes using Potri.002G079000 as the query ^19^, then the first exons of *PI*-like duplicates of them were used to construct phylogenetic tree.

### Sex-linked region specific gene identification

We identified the female and male specific genes (15W-SLR specific genes and 15Z-SLR specific genes) of the *S. polyclona* complex using BLASTP method (evalue 1e-5). The 15W-SLR specific genes means that there is no homologous gene in 15Z-SLR, and vice versa. For autotetraploid W2, we identified the 15Z(b)-SLR, 15Z(c)-SLR, and 15Z(d)-SLR specific genes, respectively. There is no homologous gene in 15Z(b)-SLR, 15Z(c)-SLR, and 15Z(d)-SLR for 15W-SLR specific genes in W2. These sex specific genes were annotated in TAIR (https://www.arabidopsis.org/). Venn diagrams were presented using EVenn ^89^.

### Gene expression analyses

We used sRNAminer v1.1.2 ^90^ to analyze and identify small RNAs of *S. polyclona* complex. Three biological replicates of buds and catkins tissues from females and males in each lineage were performed this analysis (Extended Data Table 7). SRNAanno database ^91^ was used to identify sRNAs, and adaptors, non-coding RNA including rRNA, tRNA, snoRNA, snRNA, plasmid contamination were removed from the sRNA-Seq datasets. Using sRNAminer, the clean reads were aligned to their respective reference genomes, and we applied IGV-sRNA (https://gitee.com/CJchen/IGV-sRNA) to estimate read (per site) coverage. We calculated the average read (per site) coverage of biological replicates of each partial *PI*-like duplicate and around regions.

We calculated the gene expression (excluding non-mRNA) among female and male catkins of E, E-TS, W1 and W2, respectively. Each sex and individual contained three biological replicates (Extended Data Table 7). After filtering, clean transcript reads were aligned genomes with HISAT2 ^92^, respectively, and then featureCounts ^93^ was used to calculate the number of reads mapping to each gene. Then we converted these read counts to TPM (transcripts per million reads).

## Data Availability

The genome assembly sequences have been deposited in the National Genomics Data Center (NGDC) with the BioProject number: PRJCA025570 & PRJCA016000. The sequencing datasets have been deposited in NCBI under the BioProject accession numbers PRJNA1104312.

## Acknowledgements

This study was financially supported by the National Natural Science Foundation of China (grant no. 32171813 to L.H.) and Special Fund for Scientific Research of Shanghai Landscaping & City Appearance Administrative Bureau (grant nos. G232403 & G242417 to L.H.). We are grateful to Yi Wang, Yi-Qun Liang, Qi-Chao Wu, Yan-Fei Mao, Zhi-Ying Zhu, and Guang-Nan Gong for their kind help during preparation of our study.

## Author Contributions

L.H. and Y.W. conceived, designed, and conceptualized the study. Y.W., L.H., Z.-Q. X., and R.-G. Z. performed the data analysis. L.H. collected materials. Y.W. and L.H. wrote the manuscript. J.-E. M., E.H., X.-R. W. revised the manuscript. All authors read and approved the final version of the manuscript.

