## Supplementary information for "A novel downstream factor in willows replaces the ancestral sex determining gene"

**This PDF file includes:**

**Supplementary Figures 1 to 13**

**Supplementary Tables 1 to 9**

### Supplementary Figures

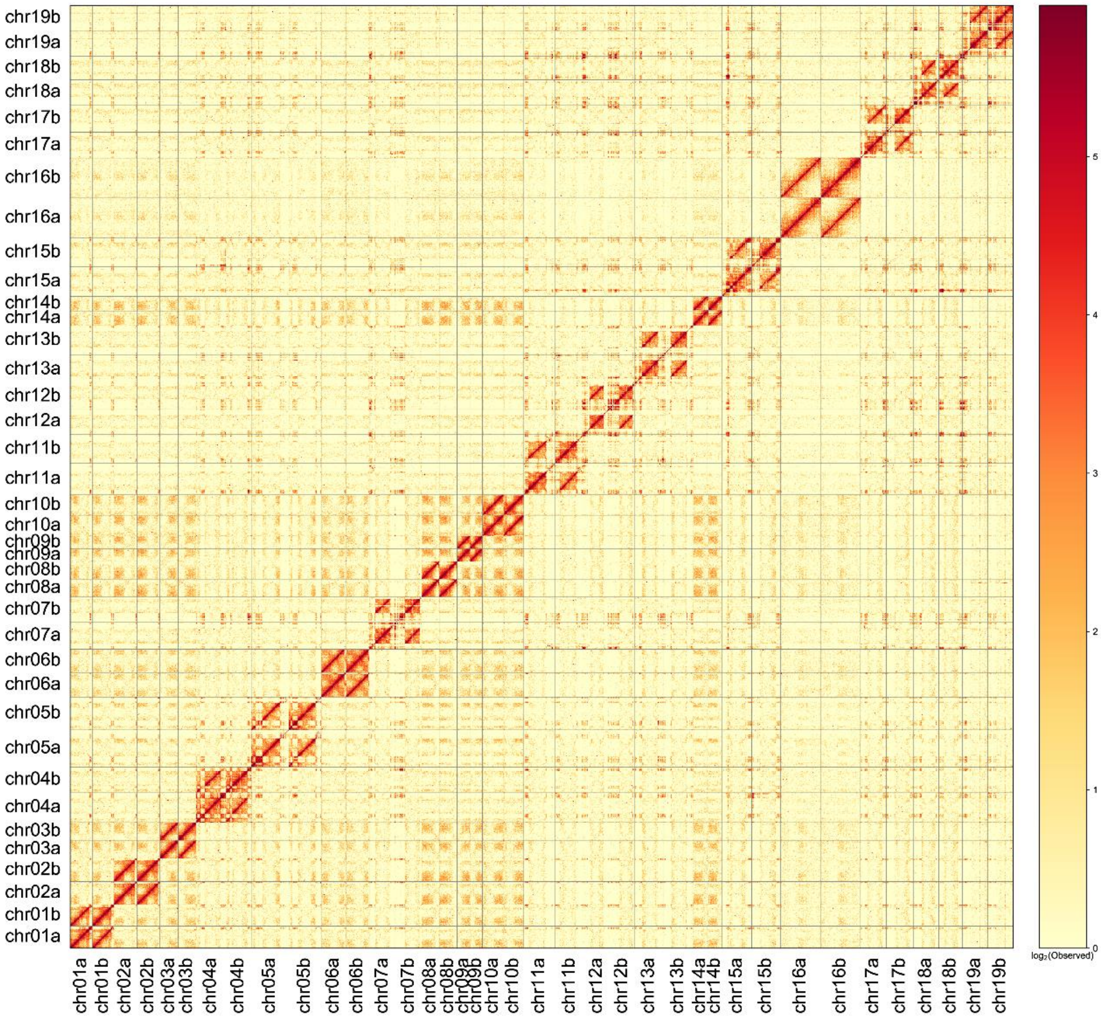


**Supplementary Fig. 1** Genome-wide analysis of chromatin interactions in the *S. polyclona*-E genome based on Hi-C data.

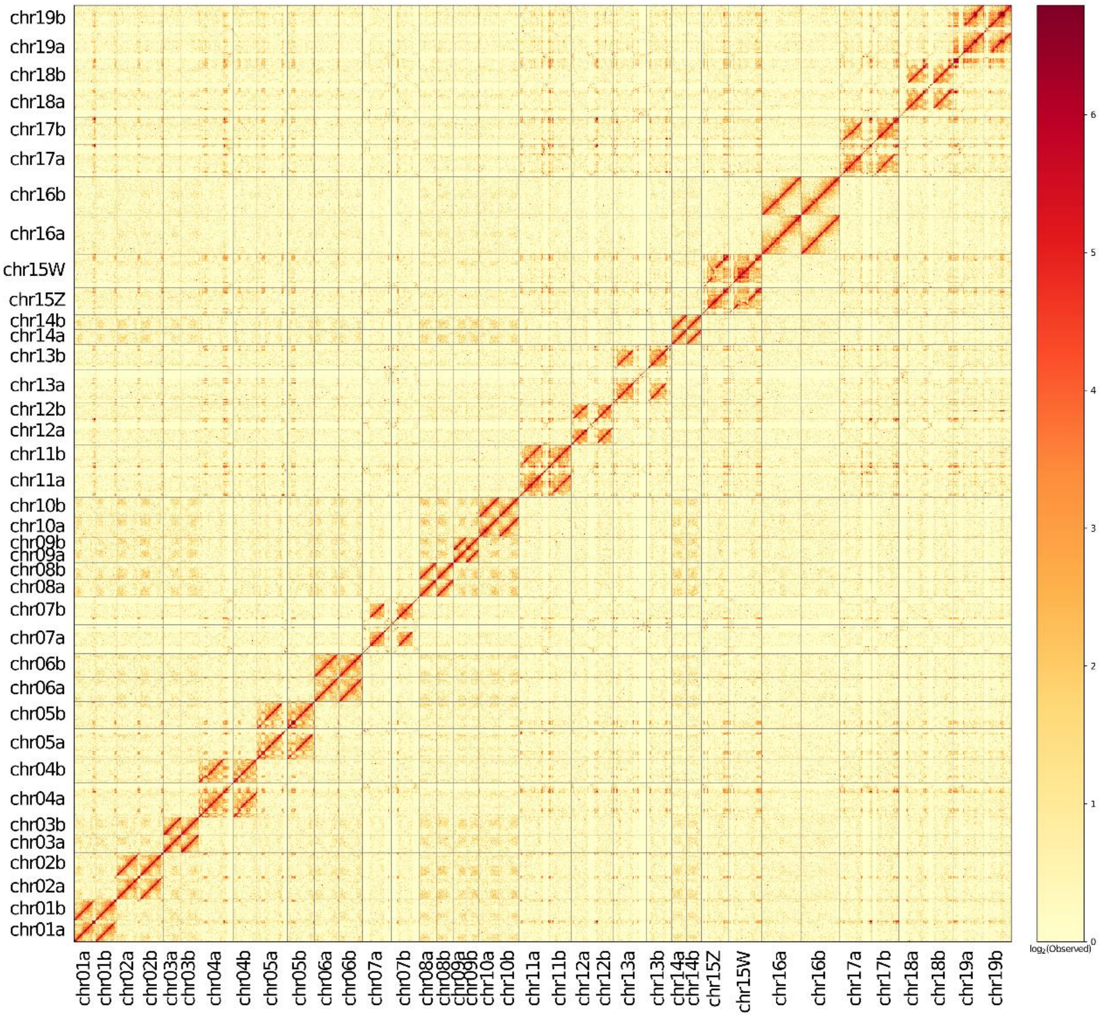


**Supplementary Fig. 2** Genome-wide analysis of chromatin interactions in the *S. polyclona*-E-TS genome based on Hi-C data.

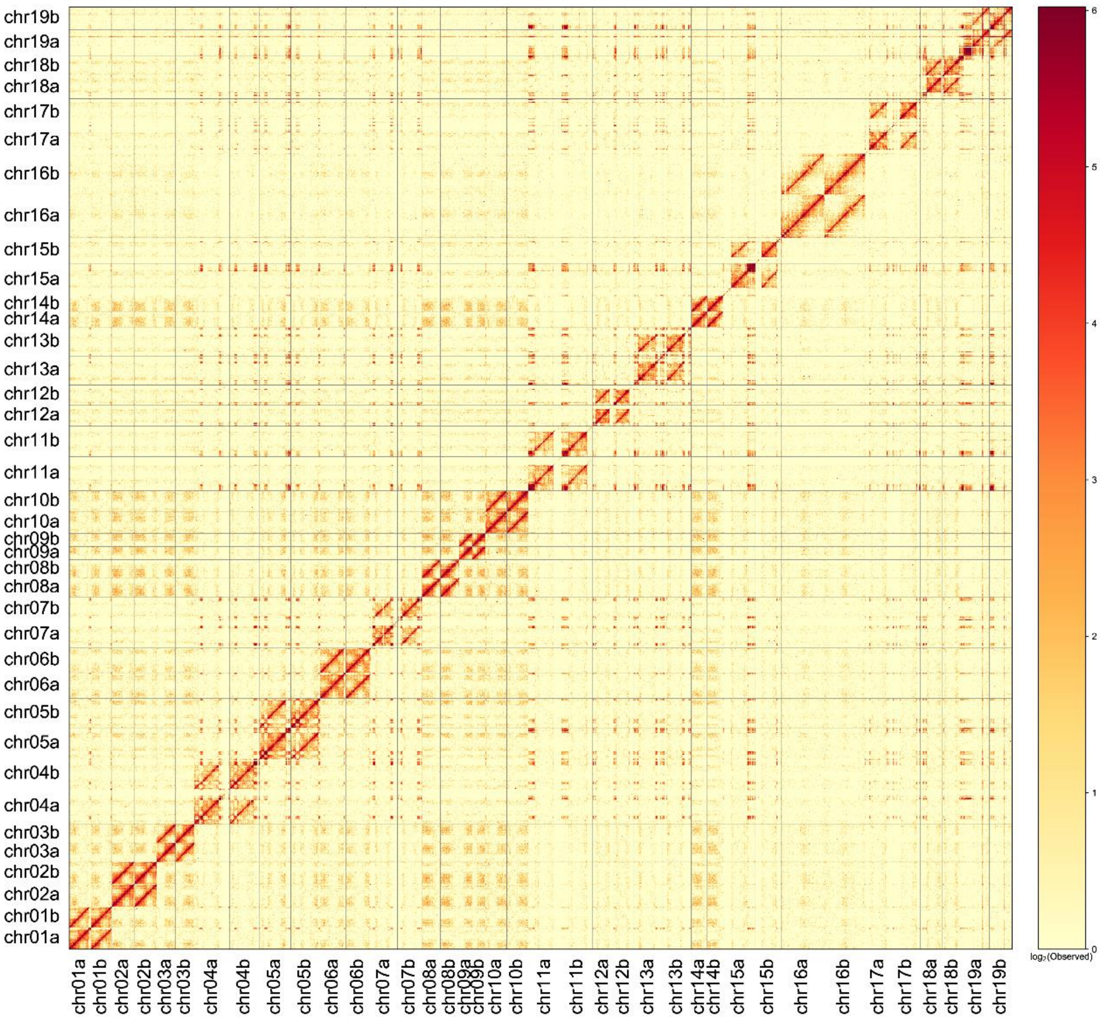


**Supplementary Fig. 3** Genome-wide analysis of chromatin interactions in the *S. polyclona*-W1 genome based on Hi-C data.

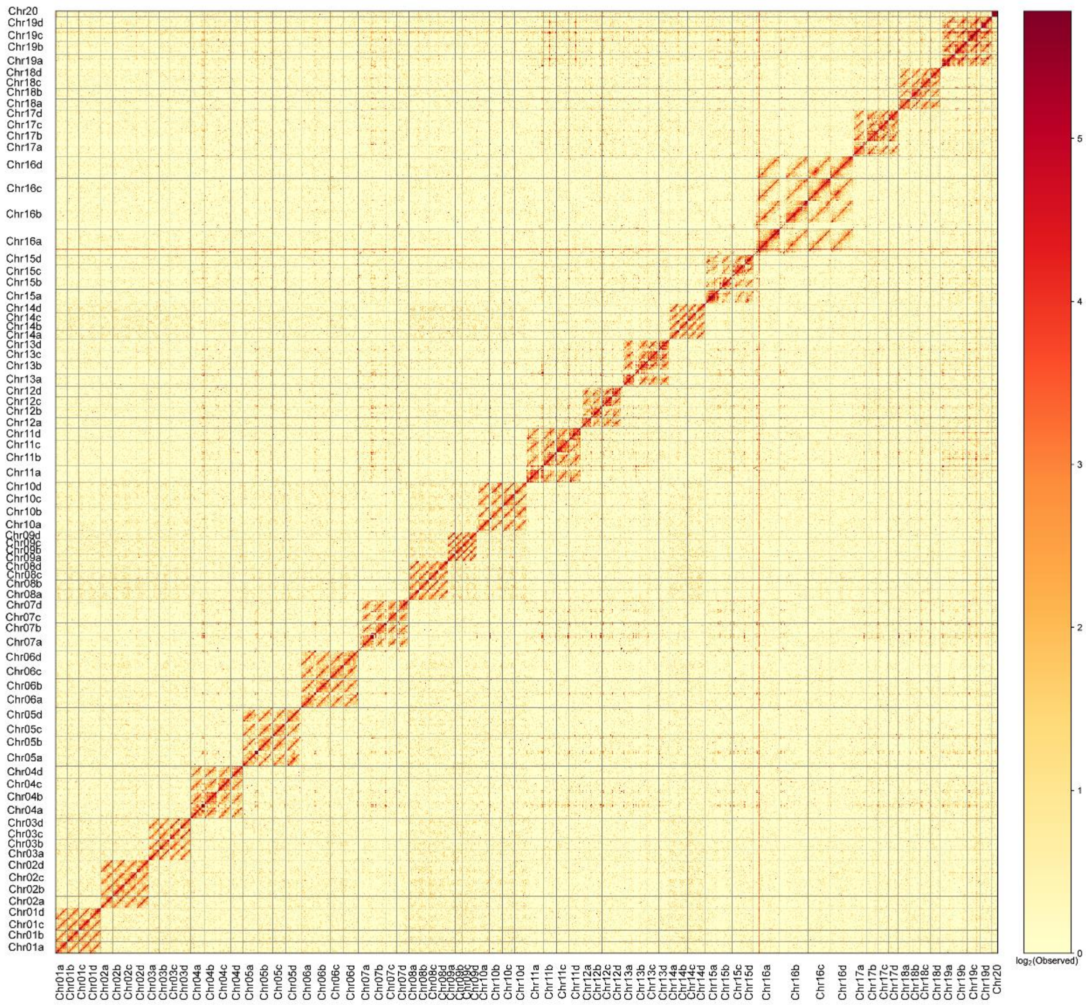


**Supplementary Fig. 4** Genome-wide analysis of chromatin interactions in the *S. polyclona*-W2 genome based on Hi-C data.

**(a)**

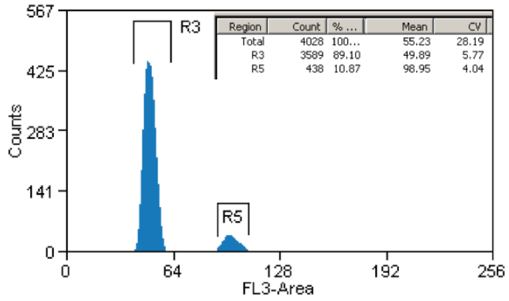


***S. polyclona* - E (HL00164)**

**(b)**

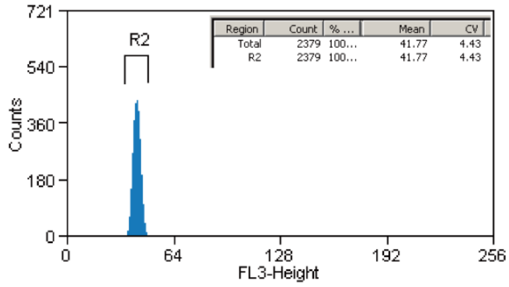


***S. polyclona* - E - TS (HL00025)**

**(c)**

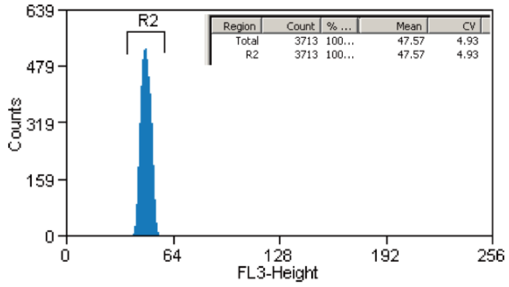


***S. polyclona* - W1 (HL00215)**

**(d)**

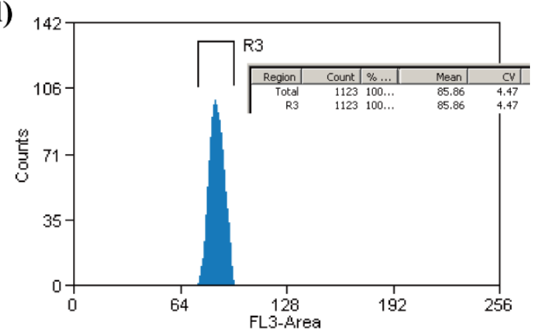


***S. polyclona* - W2 (HL00183)**

**(e)**

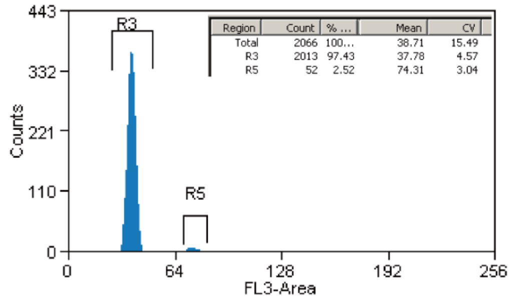


***S. baileyi***

**Supplementary Fig. 5** Flow cytometry histograms of *Salix polyclona* complex and relevant external standard. (a) *S. polyclona*-E, (b) *S. polyclona* E-TS, (c) *S. polyclona*-W1, (d) *S. polyclona*-W2, and (e) their external standard *S. baileyi*.

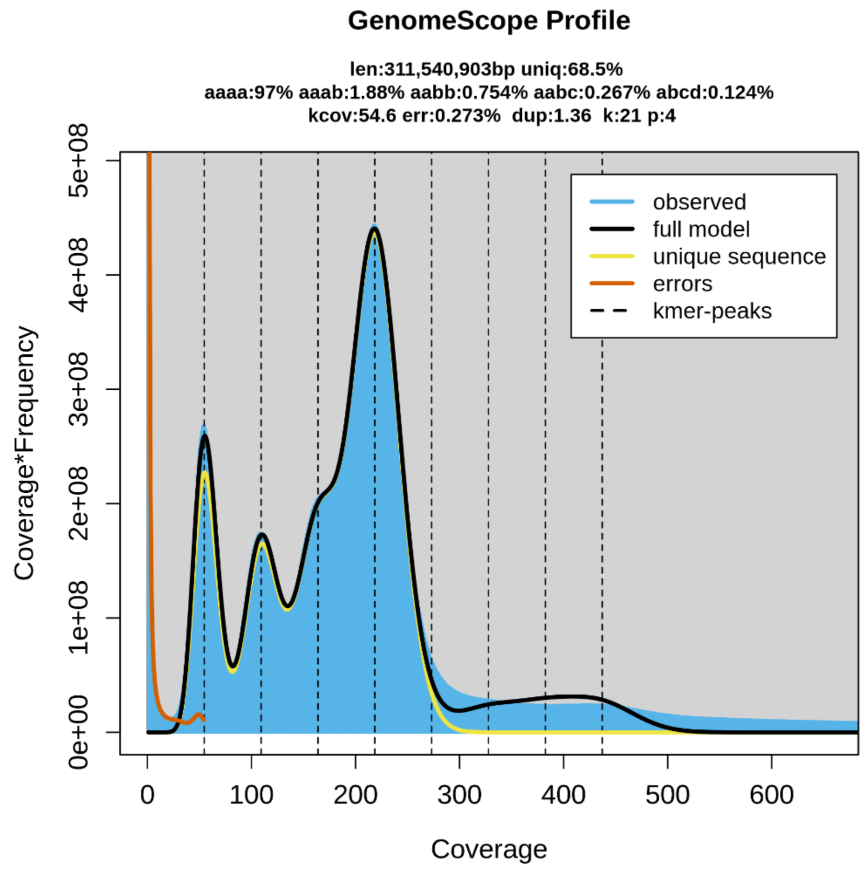


**Supplementary Fig. 6** GenomeScope plots for *S. polyclona* (HL00183). The higher proportion of aaab (1.88%) than aabb (0.754%) indicates that *S. polyclona* is autotetraploid.

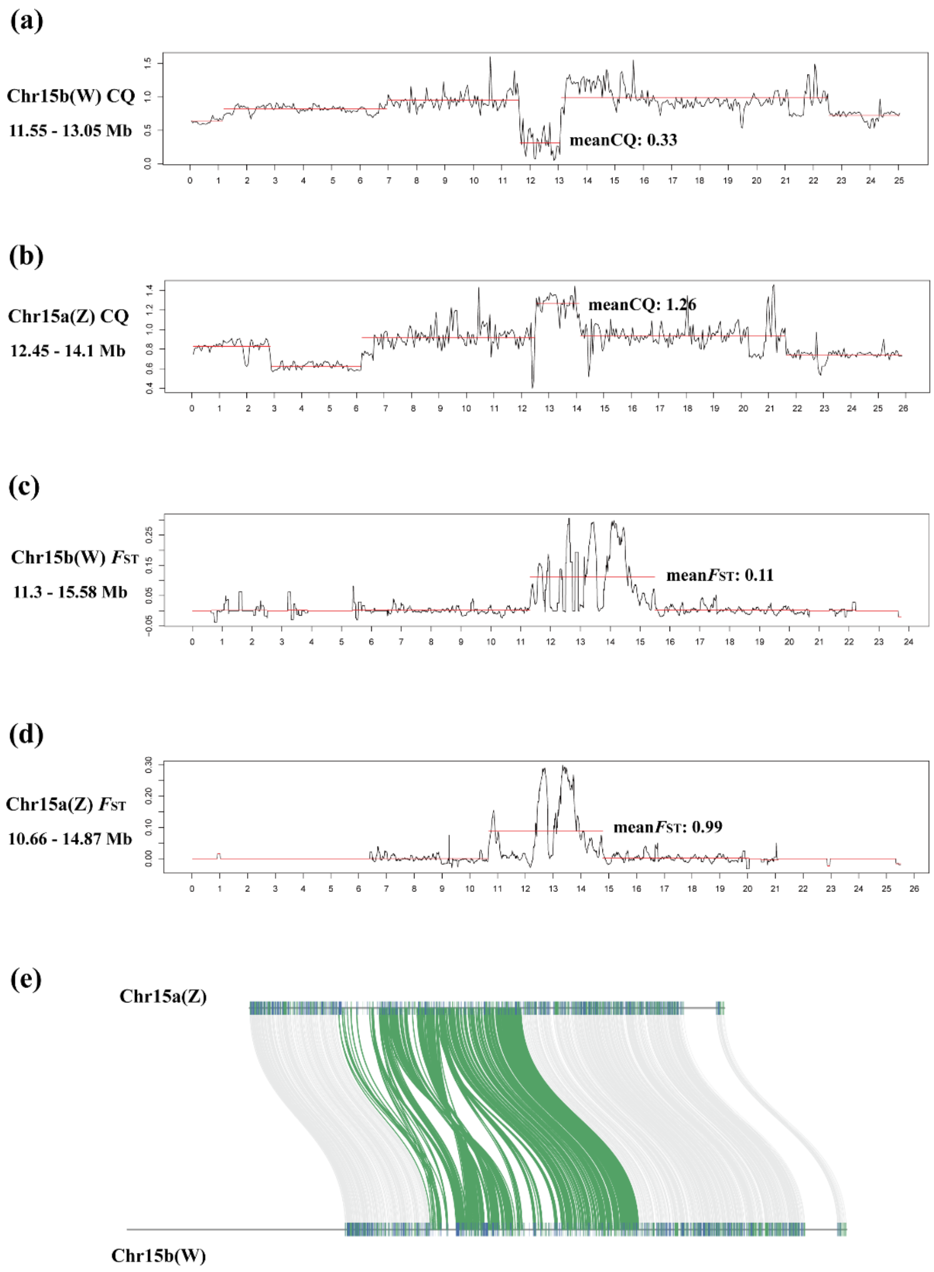


**Supplementary Fig. 7** Sex-linked region in *S. polyclona*-E. (a), (b) The CQ results in chromosome 15b and chromosome 15a. (c), (d) The $F_{ST}$ results in chromosome 15b and chromosome 15a. (e) The synteny between Z (15a) and W (15b) chromosomes.

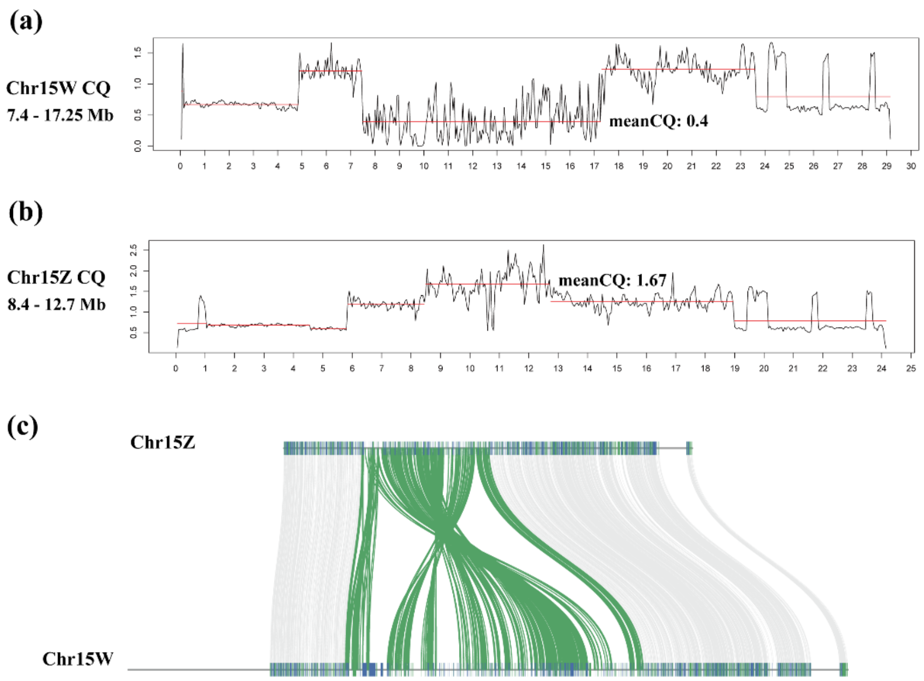


**Supplementary Fig. 8** Sex-linked region in *S. polyclona*-E-TS. (a), (b) The CQ results in chromosome 15W and chromosome 15Z. (c) The synteny between 15Z and 15W chromosomes.

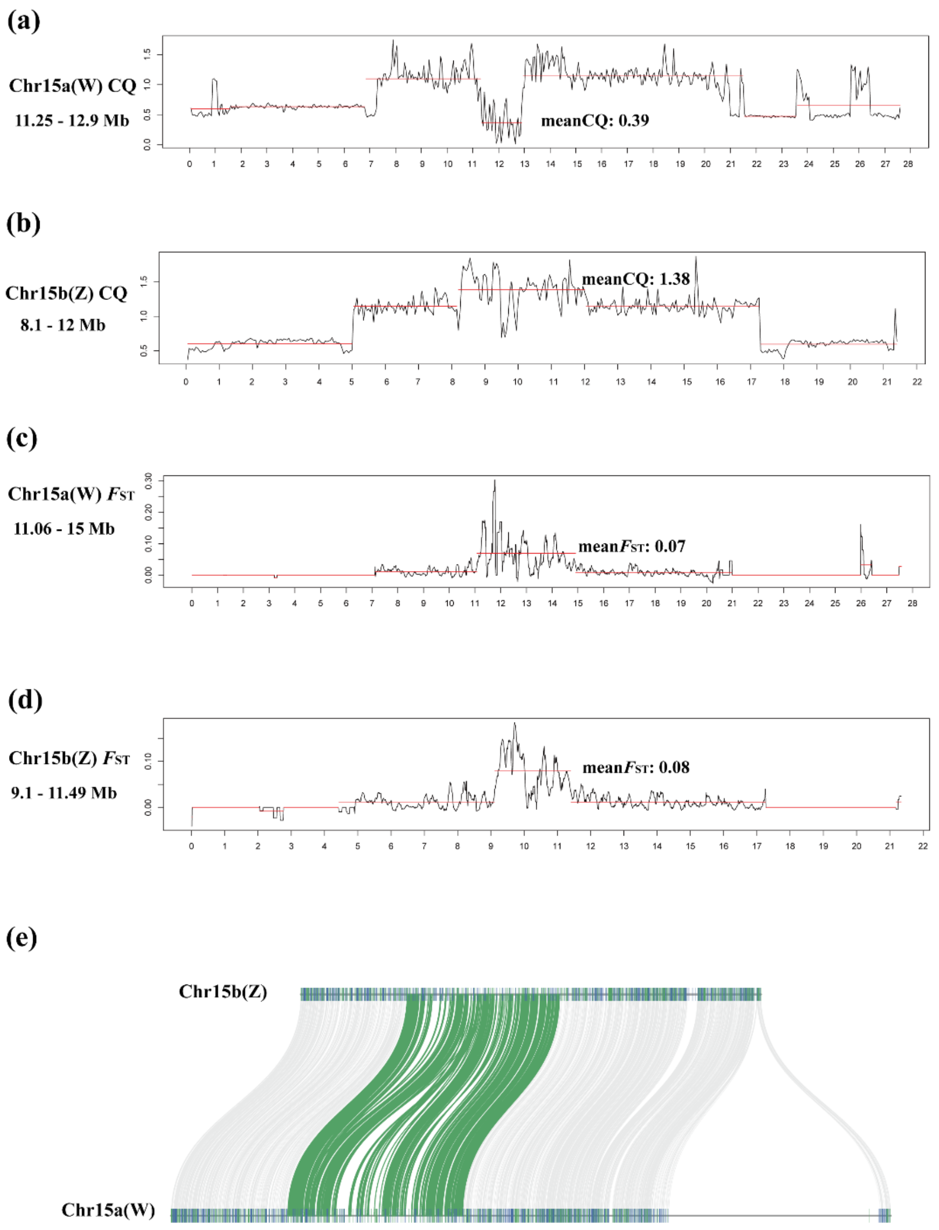


**Supplementary Fig. 9** Sex-linked region in *S. polyclona*-W1. (a), (b) The CQ results in chromosome 15a and chromosome 15b. (c), (d) The $F_{ST}$ results in chromosome 15a and chromosome 15b. (e) The synteny between Z (15b) and W (15a) chromosomes.

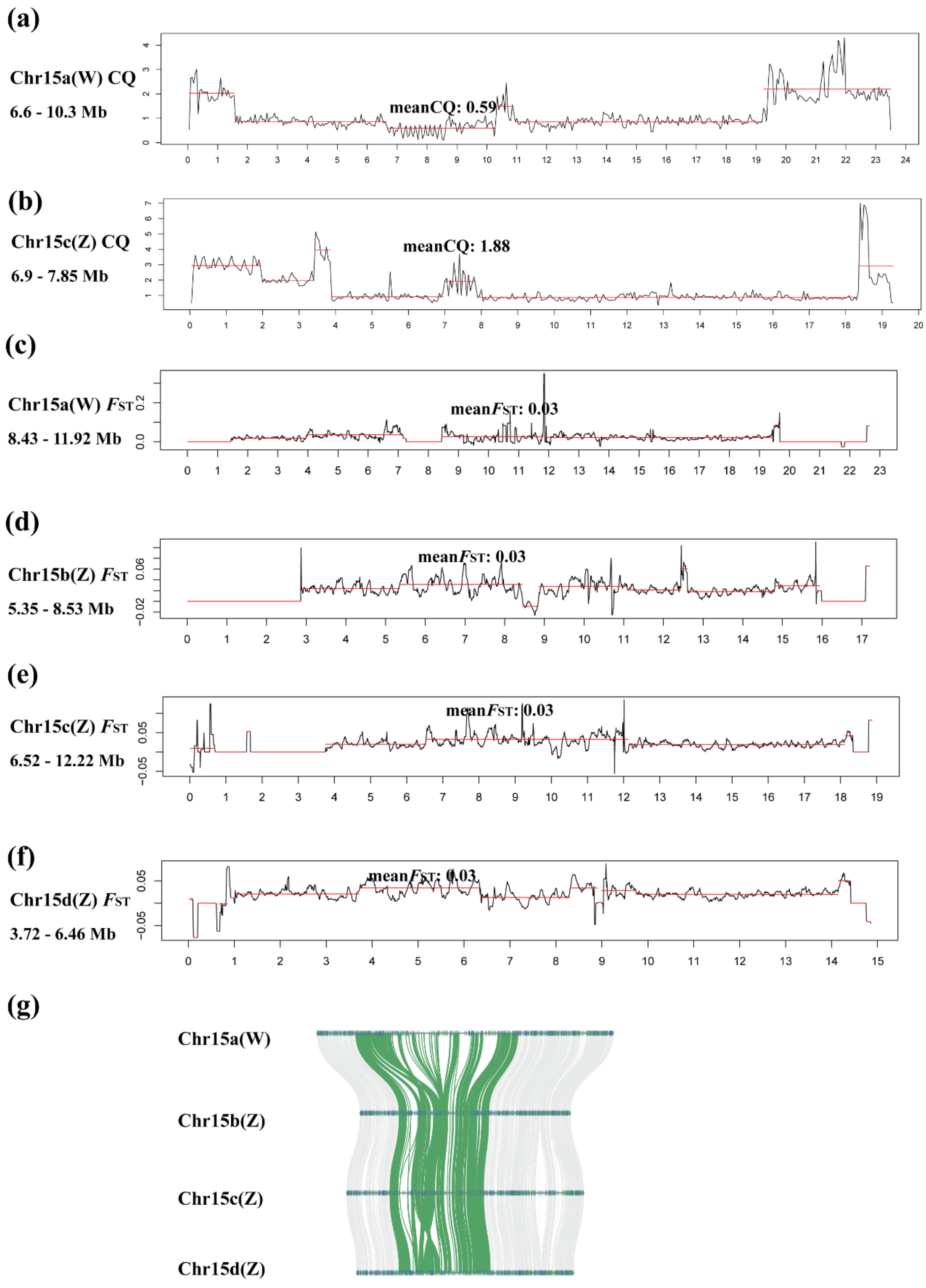


**Supplementary Fig. 10** Sex-linked region in *S. polyclona*-W2. (a) The CQ results in chromosome 15a. (b) The CQ results in chromosome 15c. (c) The $F_{ST}$ results in chromosome 15a. (d) The $F_{ST}$ results in chromosome 15b. (e) The $F_{ST}$ results in chromosome 15c. (f) The $F_{ST}$ results in chromosome 15d. (g) The synteny between Z (15b, 15c, 15d) and W (15a) chromosomes.

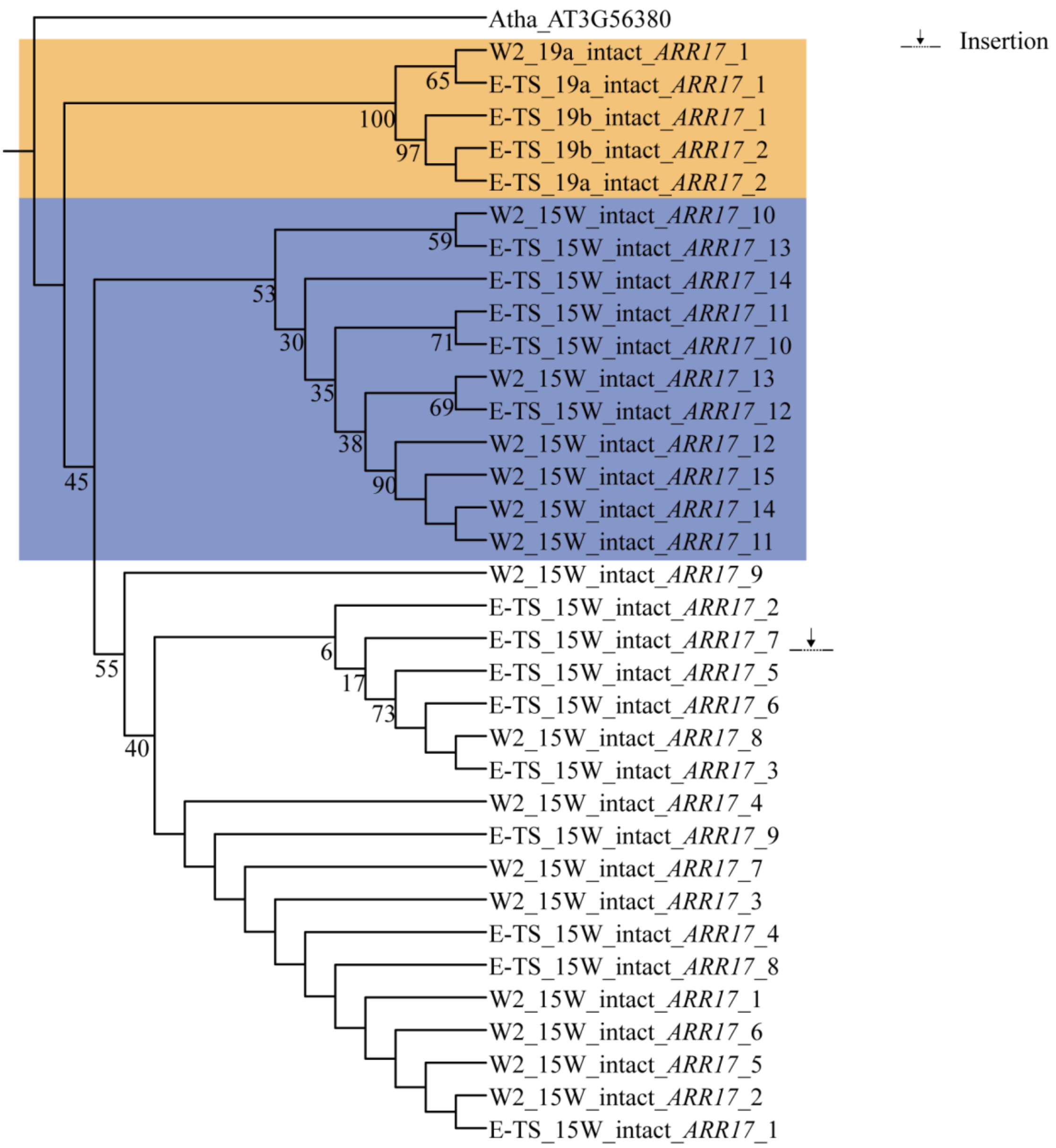


**Supplementary Fig. 11** The phylogenetic tree of intact *ARR17*-like genes in E-TS and W2. The orange and purple background color shows that the *ARR17*-like genes on chromosome 19 and 15W-SLR. The blank background color shows *ARR17*-like genes (Intact_*ARR17*_1-9) on E-TS and W2, which derived from translocation within 15W. The insertion was marked after the *ARR17*-like genes.

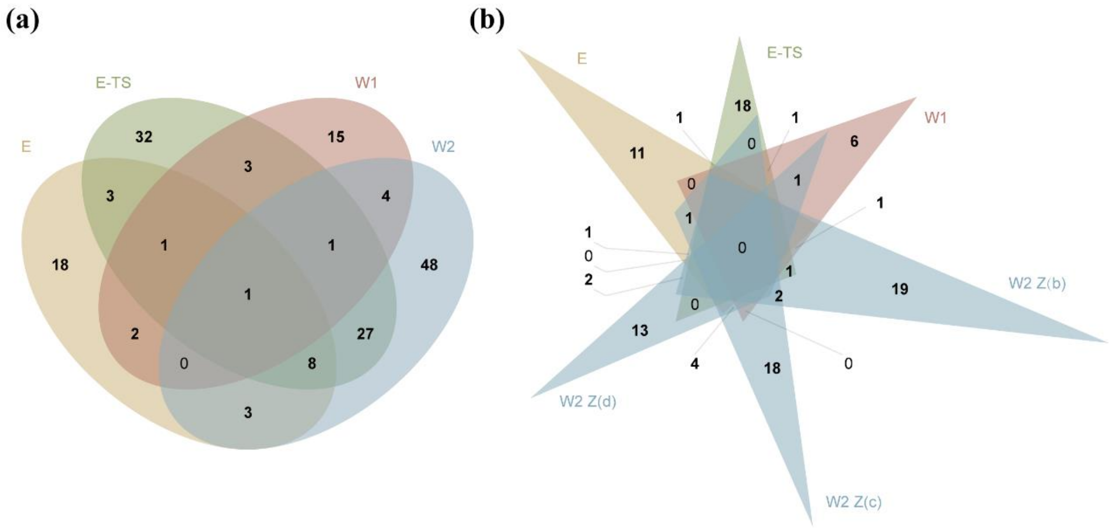


**Supplementary Fig. 12** The Venn diagram of sex specific genes. (a) The female specific genes (15W-SLR specific genes) in *S. polyclona* complex. (b) The male specific genes (15Z-SLR specific genes) in *S. polyclona* complex.

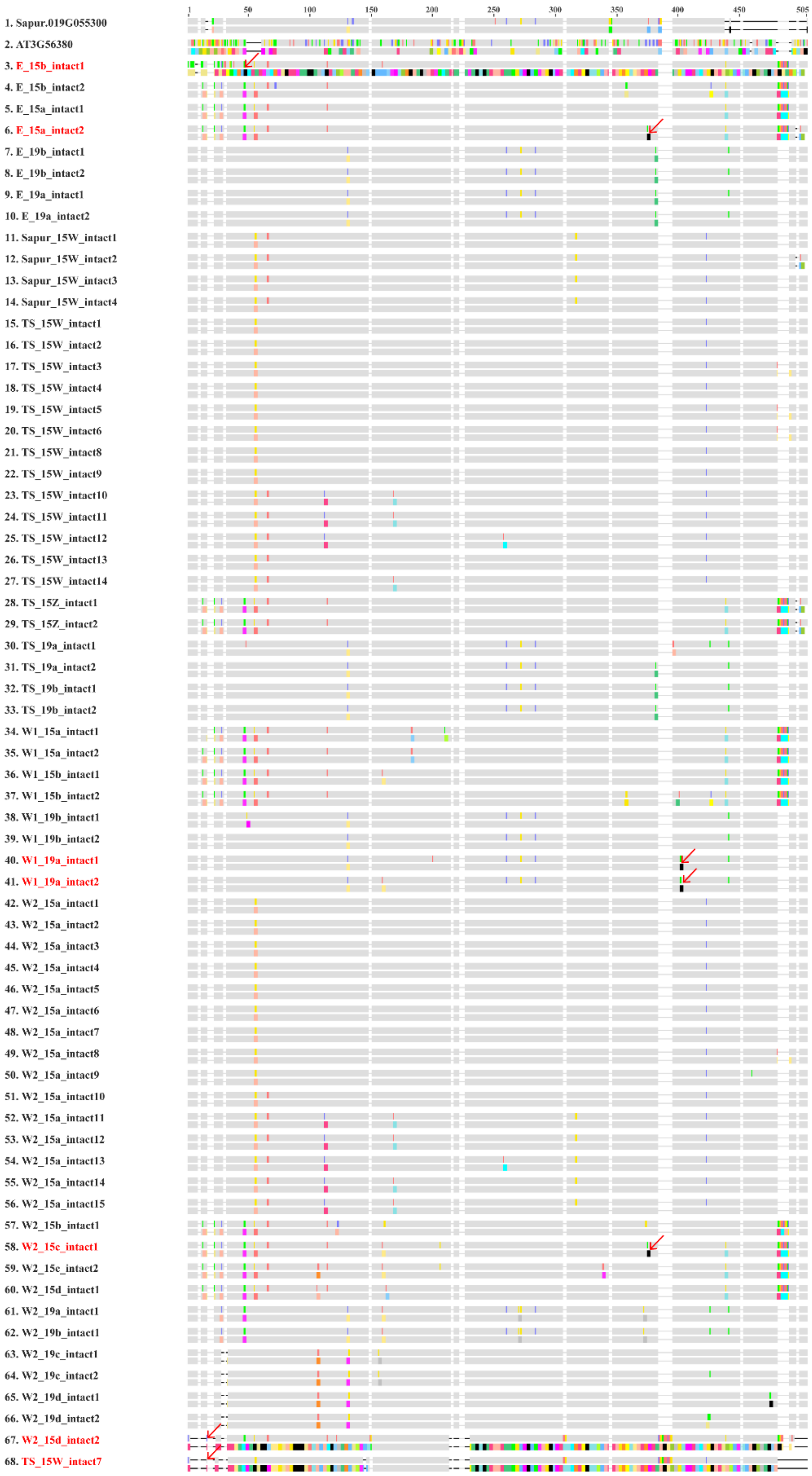


**Supplementary Fig. 13** The coding sequence and amino acid alignment result of intact *ARR17*-like duplicates in *S. polyclona* complex. The areas marked with red arrows indicate early termination or large fragment insertion.

### Supplementary Tables

**Supplementary Table 1 DNA sequencing information of *Salix polyclona* complex.**

| Lineages | Gender | Sample id | Sequencing method | Total number of reads | Total number of sequenced bases (Gb) | Average length (bp) | Depth (×) |
|---|---|---|---|---|---|---|---|
| *S. polyclona*-E | female | HL00164 | HiFi | 2,685,261 | 46.57 | 17,343 | 116 |
| *S. polyclona*-E | female | HL00164 | Illumina short reads | 278,039,000 | 41.71 | 148 | 86 |
| *S. polyclona*-E | female | HL00164 | HiC | 276,094,000 | 41.41 | 150 | 101 |
| *S. polyclona*-E-TS | female | YGDF | HiFi | 1,523,712 | 26.69 | 17,517 | 52 |
| *S. polyclona*-E-TS | female | YGDF | Illumina short reads | 233,821,000 | 35.07 | 148 | 68 |
| *S. polyclona*-E-TS | female | YGDF | HiC | 240,949,000 | 36.14 | 150 | 87 |
| *S. polyclona*-W1 | female | HL00215 | HiFi | 2,429,820 | 46.10 | 18,970 | 104 |
| *S. polyclona*-W1 | female | HL00215 | Illumina short reads | 277,311,000 | 41.60 | 148 | 82 |
| *S. polyclona*-W1 | female | HL00215 | HiC | 278,388,000 | 41.76 | 150 | 106 |
| *S. polyclona*-W2 | female | HL00183 | HiFi | 2,448,855 | 43.42 | 17,731 | 56 |
| *S. polyclona*-W2 | female | HL00183 | ONT | 1,920,267 | 54.28 | 28,269 | 72 |
| *S. polyclona*-W2 | female | HL00183 | Illumina short reads | 731,459,000 | 109.72 | 149 | 124 |
| *S. polyclona*-W2 | female | HL00183 | HiC | 685,464,000 | 102.82 | 150 | 136 |

**Supplementary Table 2 The BUSCO results of *S. polyclona* complex (protein).**

| | ***S. polyclona*-E** | ***S. polyclona*-E-TS** | ***S. polyclona*-W1** | ***S. polyclona*-W2** |
|---|---|---|---|---|
| Complete BUSCOs | 1591 (98.6%) | 1411 (98%) | 1584 (98.2%) | 1597 (98.9%) |
| Complete and single-copy BUSCOs | 50 (3.1%) | 37 (2.6%) | 56 (3.5%) | 13 (0.8%) |
| Complete and duplicated BUSCOs | 1541 (95.5%) | 1374 (95.4%) | 1528 (94.7%) | 1584 (98.1%) |
| Fragmented BUSCOs | 3 (0.2%) | 8 (0.6%) | 3 (0.2%) | 3 (0.2%) |
| Missing BUSCOs | 20 (1.2%) | 21 (1.4%) | 27 (1.6%) | 14 (0.9%) |

**Supplementary Table 3 Summary of annotated protein-coding genes in *S. polyclona* complex genomes.**

| | ***S. polyclona*-E** | ***S. polyclona*-E-TS** | ***S. polyclona*-W1** | ***S. polyclona*-W2** |
|---|---|---|---|---|
| Gene number | 62,396 | 63,175 | 62,004 | 132,186 |
| Protein coding gene number | 58,669 | 58,623 | 58,703 | 122,173 |
| Transcript number | 82,136 | 87,039 | 87,775 | 165,793 |
| Average gene region length (bp) | 3625.9 | 3614.8 | 3753.2 | 3655.5 |
| Average transcript length (bp) | 1862.2 | 1890.1 | 2066.7 | 1869.4 |
| Average coding sequence length (bp) | 1348.9 | 1343.2 | 1343.6 | 1328.1 |
| Average exons per transcript | 6.2 | 6.4 | 6.1 | 6 |
| Average exon length (bp) | 298.4 | 297.6 | 338.7 | 313.4 |
| Average intron length (bp) | 423.8 | 424.7 | 416.5 | 434.7 |

**Supplementary Table 4 Genome datasets used in the paper.**

| Species | Version | URL | References |
|---|---|---|---|
| *Salix polyclona*-E | HL00164 | this paper | |
| *Salix polyclona*-E-TS | YGDF2 | this paper | |
| *Salix polyclona*-W1 | HL00215 | this paper | |
| *Salix polyclona*-W2 | HL00183 | this paper | |
| *Salix brachista* | used in (Chen et al. 2019) | https://www.ncbi.nlm.nih.gov/assembly/GCA_009078335.1 | (Chen et al., 2019) |
| *Salix purpurea* | v5.1 | https://genome.jgi.doe.gov/portal/pages/dynamicOrganismDownload.jsf?organism=Spurpurea | (Zhou et al., 2020) |
| *Salix arbutifolia* | B649AF | https://ngdc.cncb.ac.cn/gwh/Assembly/37755/show | (Wang et al., 2020) |
| *Salix dunnii* | FNU-M-1 | https://ngdc.cncb.ac.cn/gwh/Assembly/37756/show | (He et al., 2024) |
| *Populus trichocarpa* | v4.1 | https://genome.jgi.doe.gov/portal/pages/dynamicOrganismDownload.jsf?organism=Ptrichocarpa | |

Chen, J. *et al.* Genome-wide analysis of Cushion willow provides insights into alpine plant divergence in a biodiversity hotspot. *Nat. Commun.* **10**, 5230 (2019).
Zhou, R. *et al.* A willow sex chromosome reveals convergent evolution of complex palindromic repeats. *Genome Biol.* **21**, 1–19 (2020).
Wang, Y. *et al.* Gap-free X and Y chromosomes of *Salix arbutifolia* reveal an evolutionary change from male to female heterogamety in willows, without a change in the sex-determining region. *New Phytol.* **242**, 2872–2887 (2024).
He, L. *et al.* Allopolyploidization from two dioecious ancestors leads to recurrent evolution of sex chromosomes. *Nat. Commun.* **15**, 6893 (2024).

**Supplementary Table 5 The identification information of SLRs in *S. polyclona* complex.**

| Lineage | Chr | SLR range (Mb) based on CQ results | mean CQ in the estimated SLRs | SLR range (Mb) based on *F*st results | SLR range (Mb) based on synteny analyses (Inversion) | Final SLR range based on CQ, Fst, and synteny |
|---|---|---|---|---|---|---|
| E | 15a (Z) | 12.45 - 14.10 | 1.26 | 10.66 – 14.87 | 9.29 - 10.34, 10.78 - 12.26 | 9.29 - 14.87 |
| E | 15b (W) | 11.55 - 13.05 | 0.33 | 11.30 - 15.58 | 9.24 - 9.52, 10.30 - 11.74 | 9.24 - 15.58 |
| E-TS | 15a (W) | 7.40 - 17.25 | 0.40 | / | 8.76 - 16.36 | 7.35 - 17.41 |
| E-TS | 15b (Z) | 8.40 - 12.70 | 1.67 | / | 9.09 - 12.22 | 8.40 - 12.70 |
| W1 | 15a (W) | 11.25 - 12.90 | 0.39 | 11.06 – 15.00 | 10.33 - 10.60 | 10.33 - 15.00 |
| W1 | 15b (Z) | 8.10 - 12.00 | 1.38 | 9.10 – 11.49 | 7.82 - 8.10 | 7.82 - 11.88 |
| W2 | 15a (W) | 6.60 - 10.30 | 0.59 | 8.43 - 11.92 | 4.65 - 6.54 | 3.91 - 13.82 |
| W2 | 15b (Z) | / | / | 5.35 - 8.53 | 6.80 - 7.98 | 5.35 - 10.82 |
| W2 | 15c (Z) | 6.90 - 7.85 | 1.88 | 6.52 - 12.22 | 8.03 - 9.21 | 6.48 - 12.22 |
| W2 | 15d (Z) | / | / | 3.72 - 6.46 | 4.89 - 6.17 | 3.71 - 9.28 |

**Supplementary Table 6 Distribution of protein-coding gene and RNAs on each region of the genome of *Salix polyclona* - E.**

| Chr ID | Chr length (bp) | protein-coding gene | tRNA | ncRNA | rRNA | gene |
|---|---|---|---|---|---|---|
| chr01a | 19303241 | 1810 | 34 | 17 | 0 | 1861 |
| chr01b | 18655815 | 1817 | 27 | 20 | 0 | 1864 |
| chr02a | 20202741 | 2042 | 49 | 34 | 0 | 2125 |
| chr02b | 20065287 | 2024 | 47 | 33 | 1 | 2105 |
| chr03a | 15926799 | 1665 | 36 | 21 | 0 | 1722 |
| chr03b | 15650737 | 1655 | 34 | 23 | 0 | 1712 |
| chr04a | 25928421 | 1609 | 37 | 38 | 0 | 1684 |
| chr04b | 22300902 | 1596 | 39 | 38 | 0 | 1673 |
| chr05a | 32570520 | 1927 | 50 | 68 | 0 | 2045 |
| chr05b | 28436664 | 1932 | 36 | 64 | 0 | 2032 |
| chr06a | 21309884 | 2118 | 45 | 39 | 0 | 2202 |
| chr06b | 20508203 | 2114 | 43 | 40 | 0 | 2197 |
| chr07a | 23406530 | 1177 | 19 | 28 | 2 | 1226 |
| chr07b | 22150958 | 1157 | 22 | 30 | 0 | 1209 |
| chr08a | 15581992 | 1671 | 46 | 21 | 0 | 1738 |
| chr08b | 15413492 | 1658 | 48 | 21 | 0 | 1727 |
| chr09a | 11114560 | 1325 | 42 | 17 | 2 | 1386 |
| chr09b | 11001465 | 1316 | 43 | 21 | 2 | 1382 |
| chr10a | 18038969 | 1975 | 47 | 37 | 0 | 2059 |
| chr10b | 17977600 | 1959 | 44 | 37 | 0 | 2040 |
| chr11a | 27420121 | 1170 | 32 | 27 | 7 | 1236 |
| chr11b | 25383628 | 1105 | 30 | 31 | 7 | 1173 |
| chr12a | 22334505 | 1028 | 32 | 36 | 5 | 1101 |
| chr12b | 21468393 | 1039 | 34 | 38 | 5 | 1116 |
| chr13a | 25679643 | 1199 | 26 | 42 | 29 | 1296 |
| chr13b | 25388458 | 1169 | 28 | 46 | 25 | 1268 |
| chr14a | 12898840 | 1444 | 28 | 23 | 0 | 1495 |
| chr14b | 12706730 | 1432 | 29 | 23 | 0 | 1484 |
| chr15a(Z)-SLR | 5564625 | 322 | 3 | 16 | 0 | 341 |
| chr15a(Z)-PARs | 20265916 | 930 | 20 | 25 | 0 | 975 |
| chr15b(W)-SLR | 6348526 | 380 | 4 | 17 | 0 | 401 |
| chr15b(W)-PARs | 18697762 | 916 | 23 | 28 | 0 | 967 |
| chr16a | 35036459 | 2834 | 66 | 68 | 3 | 2971 |
| chr16b | 34630559 | 2761 | 69 | 70 | 0 | 2900 |
| chr17a | 22659008 | 1125 | 20 | 17 | 2 | 1164 |
| chr17b | 23774865 | 1140 | 23 | 18 | 8 | 1189 |
| chr18a | 22056262 | 1090 | 24 | 31 | 1 | 1146 |
| chr18b | 20681245 | 1051 | 26 | 41 | 0 | 1118 |
| chr19a | 22108723 | 907 | 12 | 16 | 431 | 1366 |
| chr19b | 22040877 | 926 | 10 | 24 | 517 | 1477 |
| Pt | 155531 | 46 | 26 | 11 | 0 | 83 |
| Mt | 640961 | 108 | 20 | 12 | 0 | 140 |

**Supplementary Table 7 Distribution of protein-coding gene and RNAs on each region of the genome of *Salix polyclona* - E - TS.**

| Chr ID | Chr length (bp) | protein-coding gene | tRNA | ncRNA | rRNA | gene |
|---|---|---|---|---|---|---|
| chr01a | 19076545 | 1803 | 30 | 20 | 0 | 1853 |
| chr01b | 18554082 | 1810 | 31 | 20 | 0 | 1861 |
| chr02a | 20856483 | 2006 | 43 | 38 | 1 | 2088 |
| chr02b | 20639413 | 2005 | 49 | 38 | 1 | 2093 |
| chr03a | 15622845 | 1675 | 29 | 19 | 0 | 1723 |
| chr03b | 15586621 | 1694 | 30 | 23 | 0 | 1747 |
| chr04a | 30720841 | 1604 | 38 | 37 | 0 | 1679 |
| chr04b | 21023732 | 1609 | 36 | 40 | 0 | 1685 |
| chr05a | 26760635 | 1928 | 45 | 53 | 0 | 2026 |
| chr05b | 24107473 | 1904 | 47 | 45 | 0 | 1996 |
| chr06a | 21735010 | 2132 | 46 | 34 | 0 | 2212 |
| chr06b | 20765848 | 2089 | 46 | 34 | 0 | 2169 |
| chr07a | 25509083 | 1151 | 22 | 29 | 0 | 1202 |
| chr07b | 24637130 | 1162 | 22 | 27 | 0 | 1211 |
| chr08a | 15416969 | 1644 | 47 | 24 | 0 | 1715 |
| chr08b | 15052555 | 1629 | 44 | 22 | 0 | 1695 |
| chr09a | 11127333 | 1291 | 46 | 21 | 2 | 1360 |
| chr09b | 11068838 | 1285 | 44 | 21 | 2 | 1352 |
| chr10a | 18238504 | 1906 | 45 | 36 | 0 | 1987 |
| chr10b | 17685391 | 1894 | 42 | 31 | 0 | 1967 |
| chr11a | 27441483 | 1150 | 32 | 29 | 8 | 1219 |
| chr11b | 18868215 | 1180 | 32 | 30 | 11 | 1253 |
| chr12a | 23612362 | 975 | 30 | 34 | 2 | 1041 |
| chr12b | 13461447 | 996 | 29 | 32 | 8 | 1065 |
| chr13a | 29492733 | 1162 | 24 | 47 | 24 | 1257 |
| chr13b | 22501315 | 1169 | 27 | 43 | 24 | 1263 |
| chr14a | 13223014 | 1443 | 27 | 24 | 0 | 1494 |
| chr14b | 13102964 | 1431 | 27 | 25 | 0 | 1483 |
| chr15Z-SLR | 4296183 | 278 | 7 | 11 | 0 | 296 |
| chr15Z-PARs | 19811411 | 948 | 18 | 26 | 0 | 992 |
| chr15W-SLR | 10051440 | 502 | 6 | 20 | 0 | 528 |
| chr15W-PARs | 19069526 | 943 | 16 | 23 | 0 | 982 |
| chr16a | 34765650 | 2842 | 66 | 78 | 1 | 2987 |
| chr16b | 34322947 | 2820 | 63 | 73 | 1 | 2957 |
| chr17a | 28787173 | 1158 | 20 | 20 | 2 | 1200 |
| chr17b | 23636719 | 1126 | 21 | 17 | 2 | 1166 |
| chr18a | 25219902 | 1040 | 25 | 41 | 0 | 1106 |
| chr18b | 22530681 | 1096 | 25 | 42 | 1 | 1164 |
| chr19a | 27028549 | 1011 | 11 | 27 | 885 | 1934 |
| chr19b | 24720967 | 1007 | 9 | 21 | 926 | 1963 |
| Pt | 155503 | 34 | 25 | 13 | 0 | 72 |
| Mt | 680496 | 91 | 26 | 15 | 0 | 132 |

**Supplementary Table 8 Distribution of protein-coding gene and RNAs on each region of the genome of *Salix polyclona* - W1.**

| Chr ID | Chr length (bp) | protein-coding gene | tRNA | ncRNA | rRNA | gene |
|---|---|---|---|---|---|---|
| chr01a | 18034581 | 1832 | 35 | 18 | 0 | 1885 |
| chr01b | 17367320 | 1829 | 31 | 18 | 0 | 1878 |
| chr02a | 18923974 | 2030 | 44 | 32 | 0 | 2106 |
| chr02b | 18305473 | 2044 | 44 | 34 | 0 | 2122 |
| chr03a | 15754595 | 1677 | 32 | 21 | 0 | 1730 |
| chr03b | 15562276 | 1712 | 31 | 22 | 0 | 1765 |
| chr04a | 29467361 | 1582 | 38 | 37 | 0 | 1657 |
| chr04b | 24800913 | 1620 | 37 | 34 | 0 | 1691 |
| chr05a | 25975825 | 1917 | 43 | 53 | 0 | 2013 |
| chr05b | 24366679 | 1922 | 37 | 56 | 0 | 2015 |
| chr06a | 21399204 | 2108 | 39 | 35 | 0 | 2182 |
| chr06b | 20747234 | 2108 | 38 | 40 | 0 | 2186 |
| chr07a | 22254384 | 1151 | 24 | 25 | 0 | 1200 |
| chr07b | 20102778 | 1202 | 20 | 28 | 0 | 1250 |
| chr08a | 15514279 | 1677 | 43 | 24 | 0 | 1744 |
| chr08b | 15517819 | 1675 | 42 | 24 | 0 | 1741 |
| chr09a | 10954623 | 1313 | 39 | 20 | 2 | 1374 |
| chr09b | 10937098 | 1332 | 45 | 17 | 2 | 1396 |
| chr10a | 17816529 | 1909 | 42 | 29 | 0 | 1980 |
| chr10b | 17695820 | 1917 | 36 | 34 | 0 | 1987 |
| chr11a | 28366718 | 1129 | 30 | 26 | 8 | 1193 |
| chr11b | 25370480 | 1160 | 30 | 25 | 8 | 1223 |
| chr12a | 17697956 | 971 | 29 | 39 | 1 | 1040 |
| chr12b | 16631358 | 980 | 33 | 33 | 1 | 1047 |
| chr13a | 24035383 | 1163 | 21 | 39 | 24 | 1247 |
| chr13b | 23625313 | 1137 | 25 | 36 | 24 | 1222 |
| chr14a | 12995561 | 1453 | 24 | 24 | 0 | 1501 |
| chr14b | 12933038 | 1458 | 25 | 21 | 0 | 1504 |
| chr15a(W)-SLR | 4726261 | 297 | 3 | 14 | 0 | 314 |
| chr15a(W)-PARs | 22842617 | 983 | 18 | 26 | 0 | 1027 |
| chr15b(Z)-SLR | 4149041 | 266 | 3 | 11 | 0 | 280 |
| chr15b(Z)-PARs | 17240448 | 922 | 17 | 23 | 0 | 962 |
| chr16a | 35352969 | 2831 | 64 | 67 | 0 | 2962 |
| chr16b | 34277061 | 2785 | 73 | 63 | 1 | 2922 |
| chr17a | 22592256 | 1154 | 22 | 19 | 1 | 1196 |
| chr17b | 23484178 | 1109 | 18 | 14 | 2 | 1143 |
| chr18a | 18407715 | 1069 | 28 | 35 | 1 | 1133 |
| chr18b | 16990338 | 1106 | 32 | 29 | 1 | 1168 |
| chr19a | 22130677 | 978 | 13 | 23 | 365 | 1379 |
| chr19b | 18829412 | 1020 | 12 | 24 | 338 | 1394 |
| Pt | 155602 | 39 | 27 | 14 | 0 | 80 |
| Mt | 636913 | 136 | 20 | 9 | 0 | 165 |

**Supplementary Table 9 The statistics of haplotype genomes in *S. polyclona* complex.**

| Lineages | Average haplotype genome size | Average haplotype mRNA number |
|---|---|---|
| *S. polyclona*-E | 411.34 | 40,991 |
| *S. polyclona*-W1 | 392.09 | 43,800 |
| *S. polyclona*-E-TS | 415.07 | 43,457 |
| *S. polyclona*-W2 | 374.14 | 41,328 |
