## Supplementary figures and images for "A novel downstream factor in willows replaces the ancestral sex determining gene"

### Extended Data Fig. 1

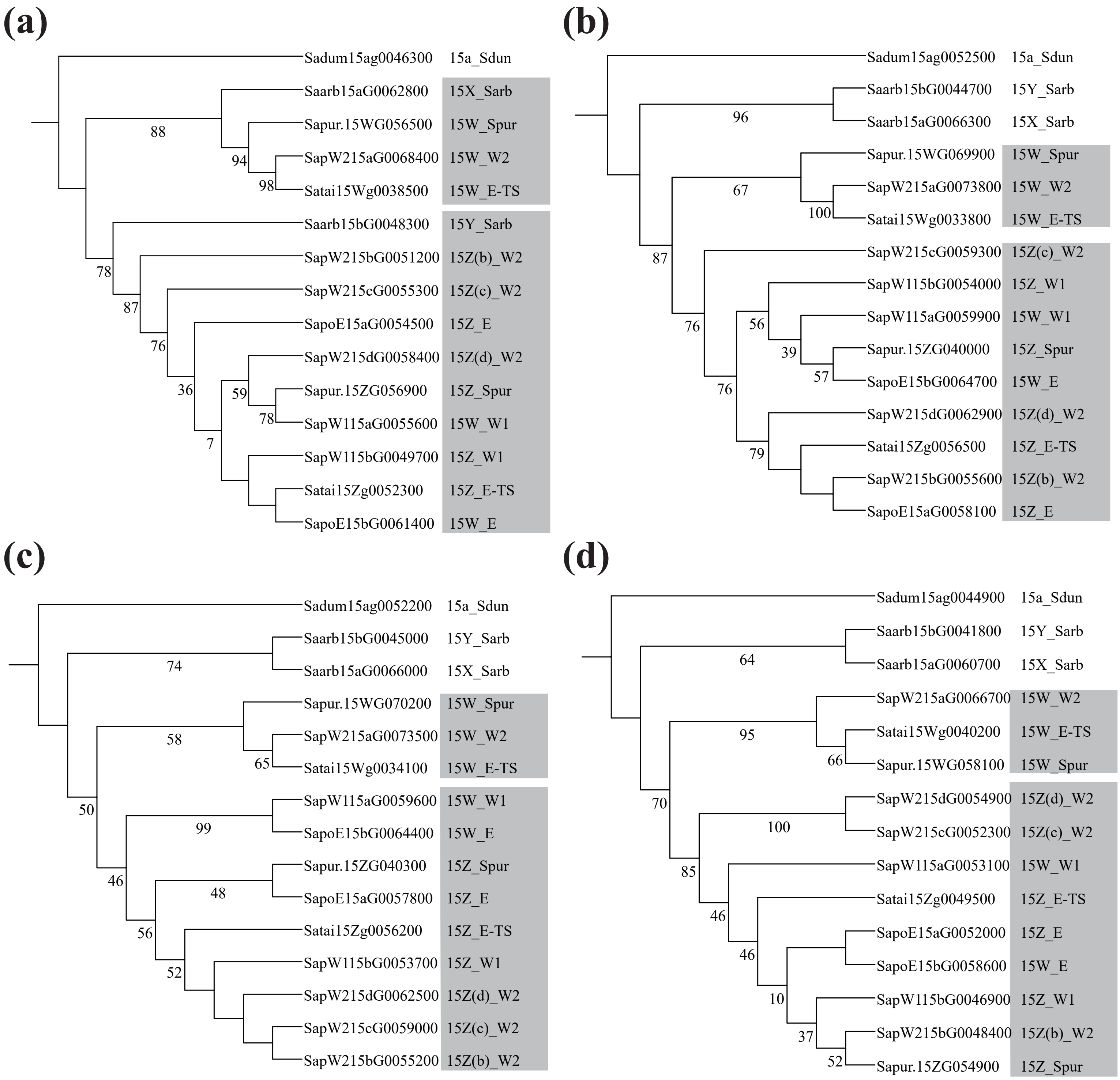
